# Genomic Evidence of Environmental and Resident *Salmonella* Senftenberg and Montevideo Contamination in the Pistachio Supply-chain

**DOI:** 10.1101/2021.03.18.436106

**Authors:** Julie Haendiges, Gordon R. Davidson, James B. Pettengill, Elizabeth Reed, Tyann Blessington, Jesse D. Miller, Nathan Anderson, Sam Myoda, Eric W. Brown, Jie Zheng, Rohan Tikekar, Maria Hoffmann

## Abstract

Pistachios have been implicated in two salmonellosis outbreaks and multiple recalls in the U.S. This study performed a retrospective data analysis of *Salmonella* associated with pistachios and a storage study to evaluate the survivability of *Salmonella* on inoculated inshell pistachios to further understand the genetics and microbiological dynamics of this commodity-pathogen pair. The retrospective data analysis on isolates associated with pistachios was performed from both short-read and long-read sequencing technologies. The sequence data were analyzed using the FDA’s Center for Food Safety and Applied Nutrition Single Nucleotide Polymorphism (SNP) analysis and Whole Genome Multi-locus Sequence Typing (wgMLST) pipeline. The storage study evaluated the survival of five strains of *Salmonella* on pistachios, both in a cocktail as well as individually. Our results demonstrate: i) evidence of persistent *Salmonella* Senftenberg and *Salmonella* Montevideo strains in pistachio environments, some of which may be due to clonal resident strains and some of which may be due to preharvest contamination; ii) presence of the Copper Homeostasis and Silver Resistance Island (CHASRI) in *Salmonella* Senftenberg and Montevideo strains in the pistachio supply chain; and iii) different serovars of *Salmonella enterica,* including *Salmonella* Senftenberg and *Salmonella* Montevideo, are able to survive in pistachios over an extended period of time.

**Importance:** Pistachios have been linked to multistate outbreaks caused by *Salmonella* serovar Senftenberg (2013, 2016) and serovar Montevideo (2016). This comprehensive study of whole-genome-sequence (WGS) data from Senftenberg and Montevideo isolates associated with pistachio outbreaks, recalls, and investigations over a nine-year period (2009-2018) was conducted to provide insights into evolutionary relationships and persistence. Multiple analytical approaches were applied including MLST, wgMLST and phylogenetic SNP analysis. The CHASRI was identified in all isolates associated with the sequence type 14 (ST14) and ST316, suggesting an adaptation in response to extrinsic pressures, such as in the farm environment. There is evidence of persistent *Salmonella* Senftenberg and Montevideo strains in pistachio pre-harvest and post-harvest environments. Utilizing metagenomic sequencing during an inoculated storage study of pistachios allowed for determination of surviving serovars over an extended period of time. Defining the mechanisms of persistence of these strains is of high importance to public health.

## Introduction

*Salmonella* has been shown to persist in low-moisture environments and represents a potential hazard for a wide range of low-moisture foods and ready-to-eat products such as nuts, peanut butter, spices, and flour (1). The ability for *Salmonella* to survive for extended periods of time in these low-moisture products and cause clinical infections is a notable public health concern (2). Contamination of these products with *Salmonella* can occur pre-harvest, during harvest, during storage, or post-harvest (e.g., after a heat-treatment step) (3).

*Salmonella enterica* subsp*. enterica* serovar Montevideo (*Salmonella* Montevideo) and *Salmonella enterica* subsp*. enterica* serovar Senftenberg (*Salmonella* Senftenberg) have been repeatedly isolated from pistachios dating back to 2008. These isolates were identified during multi-state salmonellosis outbreaks linked to pistachio consumption in 2013 and 2016 (4–7), as well as multiple product recalls across multiple time periods (e.g., 2009, 2010, 2012, 2013, 2016, and 2018) (8, 9). A previous study noted that as many as six different *Salmonella* serovars (e.g., Agona, Liverpool, Montevideo, Tennessee, Senftenberg, and Worthington) have been isolated from pistachios, and a comparison of the limited number of serovars to the numbers identified in surveys of other tree nuts may suggest a narrow and persistent contamination source (10).

Multiple potential sources of contamination with foodborne pathogens should be considered in regard to the production of produce, including pistachios. Previous orchard use should be considered in regard to total risk, for example, if the land was known to have previously been used as an animal feeding operation (11). Adjacent land use may also be a source of potential biological hazards to consider, since there is evidence that dust and microorganisms can move a short distance into adjacent tree nut orchards from an upwind animal operation (12). Moreover, pathogen-contaminated runoff can occur, and wild animals can leave waste laden with pathogens. Although many of these potential sources would concentrate on the orchard floor, there may not be any direct contact with the pistachios. However, various farming activities such as cultivation, spraying foliar treatments, and harvest also have the potential to aerosolize pathogens and deposit them on the pistachios (11).

The United States is the leading producer and exporter of pistachios in the world (13). Domestically, approximately 99% of the pistachios consumed in the United States are grown in California (14). In 2018, 264,000 bearing acres of pistachios in California resulted in a total production of 967 million pounds (15). The harvesting, transportation, wet abrasion (hulling), float tank submersion, drying, and storage of pistachios all include inherent risks for *Salmonella* contamination according to a risk assessment published in 2018 (16). The researchers were able to model atypical situations and calculate the mean risk of illness from pistachios. In particular, this study showed an increased risk of contamination due to certain pistachio production steps, including use of a float tank and a delay in drying, both of which had the greatest impact on consumer risk.

An outbreak of salmonellosis in 2016 linked to contaminated pistachios was investigated by the Centers for Disease Control and Prevention (CDC), the US Food and Drug Administration (FDA), as well as state public health labs (6). A total of eleven documented clinical cases were identified by PulseNet (nine cases of *Salmonella* Montevideo and two cases of *Salmonella* Senftenberg), with two hospitalizations in nine states. Onset dates of illnesses from patients covered a 3-month span (January to March, 2016), likely owing in part to the long shelf-life of these products. Clinical and food isolates were characterized by Pulsed Field Electrophoresis (PFGE), as well as whole genome sequencing (WGS). FDA reported raw pistachio samples collected at firms supplied by the implicated facility yielded *Salmonella* Montevideo and Senftenberg, while samples of raw pistachios collected during the investigation of the implicated facility yielded *Salmonella* Senftenberg. These *Salmonella* Senftenberg isolates were nearly indistinguishable from the clinical isolates based on WGS results (7).

Next-Generation sequencing (NGS) now regularly allows for further investigation into the genomes of *Salmonella enterica* that are responsible for illnesses, outbreaks, (17–19) and other contamination events, including a determination of whether a resident strain may be causing contamination (20). Additionally, NGS techniques provide an abundance of data regarding virulence genes, phages, and antimicrobial resistance gene patterns. The ability to capture all of these data in one assay that can be used for multiple downstream analyses makes it very appealing. FDA and CDC utilize SNP and cgMLST/wgMLST analysis in the determination of outbreaks of foodborne pathogens (21). Although these are different methods, the results are very comparable. WGS data are essential to fully characterize a bacterial pathogen of interest and can often assist in finding ways to prevent future contamination (17–19).

In this study, sequence data from *Salmonella* Senftenberg and *Salmonella* Montevideo isolates associated with pistachio outbreaks, recalls, and investigations over a nine-year period (2009-2018) were analyzed to elucidate the evolutionary relationships among these isolates and, subsequently, to better evaluate their persistence in the pistachio growing and processing environment. In addition to analyzing WGS data, in parallel we also conducted a storage experiment over a yearlong time frame that included the 2016 outbreak strains of *Salmonella* Montevideo and Senftenberg. This latter study focused on the survival rate of these strains on inshell pistachios stored at two humidities typical of low moisture foods (a_w_ < 0.7). The use of metagenomic sequencing of the cocktail-inoculated pistachios allowed for detection of the five different serovars to determine survivability of the strains over the storage period.

## Materials and methods

### Bacterial isolates used for WGS analyses

The isolates used in this study were identified by querying the Pathogen Detection tool (https://www.ncbi.nlm.nih.gov/pathogens/), a publicly available database curated by the National Center of Biotechnology Information (NCBI) (22), that routinely updates genomic sequences from food, environmental sources, and patients. SNP clusters for futher analysis were identified based on the following criteria: serovar “Montevideo” or “Senftenberg”, source “pistachio”, and location “United States”. A total of four SNP clusters (two for Senftenberg: PDS000031814, PDS000031739 and two for Montevideo: PDS000027237, PDS000032600) were identified based on our parameters. All isolates found in the SNP cluster associated with pistachios, as well as any other isolates that were closely related within the clade of the NCBI phylogenetic tree, were downloaded for analysis. The metadata (date of collection, source, and location) were also downloaded from NCBI for use in the analysis. The final data set derived from the four SNP clusters consisted of 201 isolates (106 Senftenberg and 95 Montevideo) obtained from clinical, food, and environmental sources; these were sequenced using Illumina short read technology. The sources of the 106 Senftenberg isolates consisted of almonds (1), chicken (2), clinical (7), environmental swab (43), fertilizer (1), pistachios (51), and tahini (1). These isolates were collected from 1987 to 2018. The sources of the 95 Montevideo isolates used for SNP analysis consisted of almonds (1), raw intact beef (1), cattle (1), chicken (2), clinical (30), environmental swabs (9), blend trail mix (4), missing source (2), mixed nuts (9), pistachios (35), and swine (1). These isolates were collected from 2008 to 2018. Geographically, there were 105 (Senftenberg) and 89 (Montevideo) isolates from the United States and 2 (Senftenberg) and 7 (Montevideo) from other countries (including Canada, Mexico, and the United Kingdom). The 2016 outbreak isolates were contained in SNP clusters PDS000031814 (Senftenberg) and PDS000027237 (Montevideo). There were a total of thirteen pistachio isolates (8 Senftenberg and 5 Montevideo) and 10 clinical isolates (2 Senftenberg and 8 Montevideo) associated with this outbreak that were sequenced and deposited in NCBI. The complete list of isolates and accompanying metadata can be found in Table 1. The isolates were sequenced using the Illumina sequencing chemistry by participating GenomeTrakr laboratories (23), CDC PulseNet laboratories, or private labs. Raw reads for all isolates were downloaded from the Sequence Read Archive (SRA) (https://www.ncbi.nlm.nih.gov/sra) using command line tools from the SRA toolkit available on NCBI. *De novo* assemblies were generated with SPAdes v3.13.0 (24) using k-mer lengths of [21, 33, 55, 77, 99, 127], and the options “--careful” and “--only-assembler”.

**Table 1:** Isolates and Metadata information for our study.

| Strain | Location | Isolation Source | SNP cluster | ST | Serovar | SRA | Collection Date | Facility |
| --- | --- | --- | --- | --- | --- | --- | --- | --- |
| 2016K-0180 | USA | Clinical | PDS000031814 | 14 | Senftenberg | SRR3315923 | Missing |  |
| 2017K-0050 | USA | Clinical | PDS000031814 | 14 | Senftenberg | SRR5238274 | 2016 |  |
| CFSAN000622 | USA: MD | Chicken | PDS000031814 | 14 | Senftenberg | SRR1060630 | 1987 |  |
| CFSAN015119 | USA | Pistachios | PDS000031814 | 14 | Senftenberg | SRR2585751 | 2009 |  |
| CFSAN015122 | USA | Shelled Pistachios | PDS000031814 | 14 | Senftenberg | SRR2585428 | 2009 |  |
| CFSAN016599 | USA | Pistachios | PDS000031814 | 14 | Senftenberg | SRR1816865 | 2009 | B |
| CFSAN045763 | USA:CA | Raw Pistachios | PDS000031814 | 14 | Senftenberg | SRR3168967 | 2016 | A |
| CFSAN047567 | USA:CA | Pistachios | PDS000031814 | 14 | Senftenberg | SRR3309442 | 2016 | A |
| CFSAN047866 | USA:TX | Pistachios | PDS000031814 | 14 | Senftenberg | SRR3272258 | 2016 | A |
| CFSAN048426 | USA:CA | Raw Pistachio | PDS000031814 | 14 | Senftenberg | SRR3340981 | 2016 | A |
| CFSAN048427 | USA:CA | Raw Pistachio | PDS000031814 | 14 | Senftenberg | SRR3340982 | 2016 | A |
| CFSAN048428 | USA:CA | Raw Pistachio | PDS000031814 | 14 | Senftenberg | SRR3340985 | 2016 | A |
| CFSAN048429 | USA:CA | Raw Pistachio | PDS000031814 | 14 | Senftenberg | SRR3340986 | 2016 | A |
| CFSAN058295 | USA:TX | Shelled Pistachios | PDS000031814 | 14 | Senftenberg | SRR5120752 | 2016 | A |
| FMA0102 | Canada | Pistachios | PDS000031814 | 14 | Senftenberg | SRR1258573 | 2013 |  |
| OSF033320 | USA:CA | Pistachios | PDS000031814 | 14 | Senftenberg | SRR2890018 | 2014 |  |
| OSF033321 | USA:CA | Pistachios | PDS000031814 | 14 | Senftenberg | SRR2890019 | 2014 |  |
| OSF047785 | USA:TX | Pistachios | PDS000031814 | 14 | Senftenberg | SRR3286882 | 2015 |  |
| OSF048599 | USA:CA | Pistachios | PDS000031814 | 14 | Senftenberg | SRR3372311 | 2016 |  |
| OSF048657 | USA:CA | Pistachios | PDS000031814 | 14 | Senftenberg | SRR3391910 | 2016 |  |
| OSF048658 | USA:CA | Pistachios | PDS000031814 | 14 | Senftenberg | SRR3391911 | 2016 |  |
| OSF048659 | USA:CA | Pistachios | PDS000031814 | 14 | Senftenberg | SRR3391913 | 2016 |  |
| OSF048660 | USA:CA | Pistachios | PDS000031814 | 14 | Senftenberg | SRR3657446 | 2016 |  |
| OSF048786 | USA:CA | Pistachios | PDS000031814 | 14 | Senftenberg | SRR3437480 | 2016 |  |
| OSF049331 | USA:CA | Pistachios | PDS000031814 | 14 | Senftenberg | SRR3490039 | 2016 |  |
| OSF049332 | USA:CA | Pistachios | PDS000031814 | 14 | Senftenberg | SRR3490040 | 2016 |  |
| OSF056360 | USA:CA | Pistachios | PDS000031814 | 14 | Senftenberg | SRR5057162 | 2016 |  |
| OSF056361 | USA:CA | Pistachios | PDS000031814 | 14 | Senftenberg | SRR5057161 | 2016 |  |
| OSF056362 | USA:CA | Pistachios | PDS000031814 | 14 | Senftenberg | SRR5057160 | 2016 |  |
| OSF056593 | USA:CA | Pistachios | PDS000031814 | 14 | Senftenberg | SRR5070635 | 2014 |  |
| OSF056809 | USA:CA | Raw Almonds/nuts | PDS000031814 | 14 | Senftenberg | SRR5116313 | 2014 | F |
| OSF056838 | USA:CA | Raw Pistachios | PDS000031814 | 14 | Senftenberg | SRR5120068 | 2014 |  |
| OSF058109 | USA:WA | Raw Pistachios | PDS000031814 | 14 | Senftenberg | SRR5152278 | 2014 |  |
| OSF059119 | USA:WA | Pistachios | PDS000031814 | 14 | Senftenberg | SRR5182252 | 2016 |  |
| OSF059213 | USA:CA | Environmental Sponge | PDS000031814 | 14 | Senftenberg | SRR5217564 | 2016 | G |
| OSF059253 | USA:CA | Pistachios | PDS000031814 | 14 | Senftenberg | SRR5217444 | 2016 |  |
| OSF059265 | USA:CA | Pistachios | PDS000031814 | 14 | Senftenberg | SRR5217482 | 2016 |  |
| OSF059289 | USA:WA | Pistachios | PDS000031814 | 14 | Senftenberg | SRR5229127 | 2016 |  |
| OSF059786 | USA:CA | Pistachios | PDS000031814 | 14 | Senftenberg | SRR5279048 | 2016 |  |
| OSF059815 | USA:CA | Pistachios | PDS000031814 | 14 | Senftenberg | SRR5251448 | 2016 |  |
| OSF067866 | USA:CA | Fertilizer, Bone Meal | PDS000031814 | 14 | Senftenberg | SRR5990714 | 2015 |  |
| OSF070289 | USA:CA | Environmental Sponge | PDS000031814 | 14 | Senftenberg | SRR6179039 | 2017 |  |
| OSF073486 | USA:CA | Environmental Sponge | PDS000031814 | 14 | Senftenberg | SRR6386672 | 2016 |  |
| OSF073577 | USA:CA | Pistachios | PDS000031814 | 14 | Senftenberg | SRR6424913 | 2016 |  |
| OSF074224 | USA:CA | Pistachios | PDS000031814 | 14 | Senftenberg | SRR6487131 | 2016 |  |
| OSF084195 | USA:CA | Environmental Swab | PDS000031814 | 14 | Senftenberg | SRR7643277 | 2017 | E |
| OSF090574 | USA:TX | Shelled Pistachios | PDS000031814 | 14 | Senftenberg | SRR8428217 | 2016 |  |
| OSF090575 | USA:TX | Shelled Pistachios | PDS000031814 | 14 | Senftenberg | SRR8428324 | 2016 |  |
| PNUSAS001664 | USA | Clinical | PDS000031814 | 14 | Senftenberg | SRR3240307 | 2016 |  |
| PNUSAS001739 | USA | Clinical | PDS000031814 | 14 | Senftenberg | SRR3289842 | 2016 |  |
| PNUSAS001800 | USA | Clinical | PDS000031814 | 14 | Senftenberg | SRR3392769 | Missing |  |
| PNUSAS008582 | USA | Clinical | PDS000031814 | 14 | Senftenberg | SRR5260040 | 2015 |  |
| PNUSAS140610 | USA | Clinical | PDS000031814 | 14 | Senftenberg | SRR11462365 | Missing |  |
| SGSC 2516 | USA: MD | Chicken | PDS000031814 | 14 | Senftenberg | SRR5598938 | 1987 |  |
| CFSAN017737 | USA:CA | Environmental Swab | PDS000031739 | 185 | Senftenberg | SRR3151938 | 2013 | C |
| CFSAN017738 | USA:CA | Environmental Swab | PDS000031739 | 185 | Senftenberg | SRR3151939 | 2013 | C |
| CFSAN017739 | USA:CA | Environmental Swab | PDS000031739 | 185 | Senftenberg | SRR3151940 | 2013 | C |
| CFSAN017740 | USA:CA | Environmental Swab | PDS000031739 | 185 | Senftenberg | SRR3151941 | 2013 | C |
| CFSAN017741 | USA:CA | Environmental Swab | PDS000031739 | 185 | Senftenberg | SRR3151942 | 2013 | C |
| CFSAN017744 | USA:CA | Environmental Swab | PDS000031739 | 185 | Senftenberg | SRR3151943 | 2013 | C |
| CFSAN017745 | USA:CA | Environmental Swab | PDS000031739 | 185 | Senftenberg | SRR3151944 | 2013 | C |
| CFSAN017746 | USA:CA | Environmental Swab | PDS000031739 | 185 | Senftenberg | SRR3242178 | 2013 | C |
| CFSAN017747 | USA:CA | Environmental Swab | PDS000031739 | 185 | Senftenberg | SRR3242179 | 2013 | C |
| CFSAN017748 | USA:CA | Environmental Swab | PDS000031739 | 185 | Senftenberg | SRR3242180 | 2013 | C |
| CFSAN017749 | USA:CA | Environmental Swab | PDS000031739 | 185 | Senftenberg | SRR3242181 | 2013 | C |
| CFSAN017750 | USA:CA | Environmental Swab | PDS000031739 | 185 | Senftenberg | SRR3242182 | 2013 | C |
| CFSAN017751 | USA:CA | Environmental Swab | PDS000031739 | 185 | Senftenberg | SRR3242184 | 2013 | C |
| CFSAN017752 | USA:CA | Environmental Swab | PDS000031739 | 185 | Senftenberg | SRR3242185 | 2013 | C |
| CFSAN017753 | USA:CA | Environmental Swab | PDS000031739 | 185 | Senftenberg | SRR3340442 | 2013 | C |
| CFSAN017754 | USA:CA | Environmental Swab | PDS000031739 | 185 | Senftenberg | SRR3340443 | 2013 | C |
| CFSAN032229 | USA:CA | Environmental Swab | PDS000031739 | 185 | Senftenberg | SRR1979272 | 2015 | C |
| CFSAN032230 | USA:CA | Environmental Swab | PDS000031739 | 185 | Senftenberg | SRR1982146 | 2015 | C |
| CFSAN032231 | USA:CA | Environmental Swab | PDS000031739 | 185 | Senftenberg | SRR1979316 | 2015 | C |
| CFSAN086744 | USA:PA | Pistachios | PDS000031739 | 185 | Senftenberg | SRR8113236 | 2018 | C |
| CFSAN087304 | USA:NJ | Pistachios | PDS000031739 | 185 | Senftenberg | SRR8261878 | 2018 | C |
| FNW19i50 | USA:CA | Pistachios | PDS000031739 | 185 | Senftenberg | SRR1614998 | 2013 | C |
| FNW19i51 | USA:CA | Pistachios | PDS000031739 | 185 | Senftenberg | SRR1646543 | 2013 | C |
| FSE0085 | Mexico | Tahini | PDS000031739 | 185 | Senftenberg | SRR1212317 | 2011 |  |
| FSF0102 | USA:CA | Environmental Swab | PDS000031739 | 185 | Senftenberg | SRR1535718 | 2013 | C |
| FSW0103 | USA:CA | Dry Roasted Pistachios | PDS000031739 | 185 | Senftenberg | SRR1257291 | 2013 | C |
| FSW0104 | USA:CA | Dry Roasted Pistachios | PDS000031739 | 185 | Senftenberg | SRR1257282 | 2013 | C |
| OSF056673 | USA:CA | Environmental Swab | PDS000031739 | 185 | Senftenberg | SRR5099892 | 2014 | C |
| OSF056674 | USA:CA | Environmental Swab | PDS000031739 | 185 | Senftenberg | SRR5099891 | 2014 | C |
| OSF056675 | USA:CA | Environmental Swab | PDS000031739 | 185 | Senftenberg | SRR5099888 | 2014 | C |
| OSF056676 | USA:CA | Environmental Swab | PDS000031739 | 185 | Senftenberg | SRR5099890 | 2014 | C |
| OSF056753 | USA:CA | Environmental Swab | PDS000031739 | 185 | Senftenberg | SRR5105418 | 2014 | C |
| OSF056767 | USA:CA | Environmental Swab | PDS000031739 | 185 | Senftenberg | SRR5116302 | 2014 | C |
| OSF056768 | USA:CA | Environmental Swab | PDS000031739 | 185 | Senftenberg | SRR5116309 | 2014 | C |
| OSF056769 | USA:CA | Environmental Swab | PDS000031739 | 185 | Senftenberg | SRR5116457 | 2014 | C |
| OSF057938 | USA:CA | Environmental Swab | PDS000031739 | 185 | Senftenberg | SRR5120074 | 2014 | D |
| OSF057939 | USA:CA | Environmental Swab | PDS000031739 | 185 | Senftenberg | SRR5120108 | 2014 | D |
| OSF057940 | USA:CA | Environmental Swab | PDS000031739 | 185 | Senftenberg | SRR5120103 | 2014 | D |
| OSF057941 | USA:CA | Environmental Swab | PDS000031739 | 185 | Senftenberg | SRR5120104 | 2014 | D |
| OSF057942 | USA:CA | Environmental Swab | PDS000031739 | 185 | Senftenberg | SRR5120105 | 2014 | D |
| OSF057943 | USA:CA | Environmental Swab | PDS000031739 | 185 | Senftenberg | SRR5120107 | 2014 | D |
| OSF057945 | USA:CA | Environmental Swab | PDS000031739 | 185 | Senftenberg | SRR5120102 | 2014 | D |
| OSF057946 | USA:CA | Environmental Swab | PDS000031739 | 185 | Senftenberg | SRR5230332 | 2014 | D |
| OSF057947 | USA:CA | Environmental Swab | PDS000031739 | 185 | Senftenberg | SRR5230170 | 2014 | D |
| OSF057948 | USA:CA | Environmental Swab | PDS000031739 | 185 | Senftenberg | SRR5230325 | 2014 | D |
| OSF057951 | USA:CA | Environmental Swab | PDS000031739 | 185 | Senftenberg | SRR5230171 | 2014 | D |
| OSF057978 | USA:CA | Raw Pistachios | PDS000031739 | 185 | Senftenberg | SRR5230465 | 2014 | C |
| OSF057979 | USA:CA | Raw Pistachios | PDS000031739 | 185 | Senftenberg | SRR5230320 | 2014 | C |
| OSF057980 | USA:CA | Raw Pistachios | PDS000031739 | 185 | Senftenberg | SRR5230330 | 2014 | C |
| OSF057981 | USA:CA | Raw Pistachios | PDS000031739 | 185 | Senftenberg | SRR5230168 | 2014 | C |
| OSF057982 | USA:CA | Raw Pistachios | PDS000031739 | 185 | Senftenberg | SRR5230385 | 2014 | C |
| OSF087751 | USA:CA | Finished Product Pistachios | PDS000031739 | 185 | Senftenberg | SRR8176592 | 2018 | C |
| 44713 | United Kingdom | Clinical | PDS000027237 | 316 | Montevideo | SRR1957791 | 2014 |  |
| 378603 | United Kingdom | Clinical | PDS000027237 | 316 | Montevideo | SRR8568765 | 2017 |  |
| 533421 | United Kingdom | Clinical | PDS000027237 | 316 | Montevideo | SRR8515722 | 2018 |  |
| 570824 | United Kingdom | Clinical | PDS000027237 | 316 | Montevideo | SRR8484198 | 2018 |  |
| 708223 | United Kingdom | Clinical | PDS000027237 | 316 | Montevideo | SRR8774349 | 2019 |  |
| 315731156 | Missing | Pistachios | PDS000027237 | 316 | Montevideo | SRR500745 | 2009 | B |
| 2016K-0167 | USA | Clinical | PDS000027237 | 316 | Montevideo | SRR3278065 | Missing |  |
| CFSAN000244 | Missing | Pistachios | PDS000027237 | 316 | Montevideo | SRR500472 | 2009 | B |
| CFSAN013604 | USA | Pistachios | PDS000027237 | 316 | Montevideo | SRR2559381 | 2008 |  |
| CFSAN014882 | USA | Pistachios | PDS000027237 | 316 | Montevideo | SRR1973753 | 2009 |  |
| CFSAN015113 | USA | Blend Trail Mix | PDS000027237 | 316 | Montevideo | SRR2559383 | 2008 | B |
| CFSAN015114 | USA | Blend Trail Mix | PDS000027237 | 316 | Montevideo | SRR2586832 | 2008 | B |
| CFSAN015115 | USA | Pistachios | PDS000027237 | 316 | Montevideo | SRR2585772 | 2009 | B |
| CFSAN015116 | USA | Blend Trail Mix | PDS000027237 | 316 | Montevideo | SRR2586844 | 2009 | B |
| CFSAN015117 | USA | cashew/almond/pistachio blend | PDS000027237 | 316 | Montevideo | SRR2585430 | 2009 | B |
| CFSAN015118 | USA | Pistachios | PDS000027237 | 316 | Montevideo | SRR2585429 | 2009 | B |
| CFSAN015120 | USA | Pistachios | PDS000027237 | 316 | Montevideo | SRR2585750 | 2009 | B |
| CFSAN015123 | USA | Blend Trail Mix | PDS000027237 | 316 | Montevideo | SRR2559382 | 2008 | B |
| CFSAN016379 | USA:CA | Environmental Swab | PDS000027237 | 316 | Montevideo | SRR1816812 | 2009 | B |
| CFSAN016380 | USA:CA | roasted & salted garlic/onion pistachios | PDS000027237 | 316 | Montevideo | SRR1816860 | 2009 | B |
| CFSAN016411 | USA:CA | Environmental Swab | PDS000027237 | 316 | Montevideo | SRR1816817 | 2009 | B |
| CFSAN016412 | USA:CA | Environmental Swab | PDS000027237 | 316 | Montevideo | SRR1816831 | 2009 | B |
| CFSAN016600 | USA | Pistachios | PDS000027237 | 316 | Montevideo | SRR1816838 | 2009 | B |
| CFSAN016601 | USA | Pistachios | PDS000027237 | 316 | Montevideo | SRR1816839 | 2009 | B |
| CFSAN037640 | USA:IL | Shell Pistachios | PDS000027237 | 316 | Montevideo | SRR3038647 | 2015 | A |
| CFSAN045764 | USA:CA | Raw Pistachios | PDS000027237 | 316 | Montevideo | SRR3168966 | 2016 | A |
| CFSAN047166 | USA:CA | Pistachios | PDS000027237 | 316 | Montevideo | SRR3242198 | 2016 | A |
| CFSAN047167 | USA:CA | Pistachios | PDS000027237 | 316 | Montevideo | SRR3242197 | 2016 | A |
| CFSAN047168 | USA:CA | Pistachios | PDS000027237 | 316 | Montevideo | SRR3242196 | 2016 | A |
| CFSAN047169 | USA:CA | Pistachios | PDS000027237 | 316 | Montevideo | SRR3242195 | 2016 | A |
| CFSAN051296 | USA | Raw Pistachios | PDS000027237 | 316 | Montevideo | SRR3707433 | 2016 |  |
| CFSAN064799 | USA | Shelled Pistachios | PDS000027237 | 316 | Montevideo | SRR5758423 | 2017 | K |
| FSIS1606641 | USA:MI | Swine | PDS000027237 | 316 | Montevideo | SRR3555186 | 2016 |  |
| FSIS1709808 | USA:GA | Raw Intact Chicken | PDS000027237 | 316 | Montevideo | SRR5196071 | 2016 |  |
| FSL_R8-4916 | Missing | Missing | PDS000027237 | 316 | Montevideo | SRR493332 | Missing |  |
| FSL_R8-4922 | Missing | Missing | PDS000027237 | 316 | Montevideo | SRR493336 | Missing |  |
| OSF045712 | USA:IN | Chicken | PDS000027237 | 316 | Montevideo | SRR3191464 | 2015 | H |
| OSF046987 | USA:CA | Nuts | PDS000027237 | 316 | Montevideo | SRR3225384 | 2015 | F |
| OSF047721 | USA:CA | Pistachios | PDS000027237 | 316 | Montevideo | SRR3294561 | 2015 |  |
| OSF051202 | USA:WA | Pistachios | PDS000027237 | 316 | Montevideo | SRR3606932 | 2016 |  |
| OSF056837 | USA:CA | Raw Pistachios | PDS000027237 | 316 | Montevideo | SRR5120069 | 2014 |  |
| OSF059120 | USA:WA | Pistachios | PDS000027237 | 316 | Montevideo | SRR5182248 | 2016 |  |
| OSF069055 | USA:CA | Environmental sponge | PDS000027237 | 316 | Montevideo | SRR6108679 | 2015 |  |
| OSF069128 | USA:CA | Raw Pistachios | PDS000027237 | 316 | Montevideo | SRR6109685 | 2015 |  |
| OSF075603 | USA:CA | Pistachios | PDS000027237 | 316 | Montevideo | SRR6848181 | 2016 |  |
| OSF075630 | USA:WA | Pistachios | PDS000027237 | 316 | Montevideo | SRR6782992 | 2016 |  |
| OSF075637 | USA:CA | Pistachios | PDS000027237 | 316 | Montevideo | SRR6848879 | 2016 |  |
| OSF075638 | USA:CA | Pistachios | PDS000027237 | 316 | Montevideo | SRR6848843 | 2016 |  |
| OSF076741 | USA:CA | Pistachios | PDS000027237 | 316 | Montevideo | SRR6848188 | 2016 |  |
| OSF076742 | USA:CA | Pistachios | PDS000027237 | 316 | Montevideo | SRR6848319 | 2016 |  |
| OSF076748 | USA:CA | Pistachios | PDS000027237 | 316 | Montevideo | SRR6849216 | 2016 |  |
| OSF076749 | USA:CA | Pistachios | PDS000027237 | 316 | Montevideo | SRR6849236 | 2016 |  |
| OSF076750 | USA:CA | Pistachios | PDS000027237 | 316 | Montevideo | SRR6849235 | 2016 | G |
| OSF077989 | USA:CA | Environmental sponge | PDS000027237 | 316 | Montevideo | SRR6971655 | 2016 |  |
| OSF077990 | USA:CA | Environmental sponge | PDS000027237 | 316 | Montevideo | SRR6967965 | 2016 |  |
| OSF078698 | USA:CA | Environmental sponge | PDS000027237 | 316 | Montevideo | SRR7064464 | 2016 |  |
| OSF084050 | USA:CA | Finished Almonds | PDS000027237 | 316 | Montevideo | SRR7638853 | 2017 | J |
| OSF088275 | USA:CA | Nuts | PDS000027237 | 316 | Montevideo | SRR8272032 | 2018 | G |
| OSF088276 | USA:CA | Nuts | PDS000027237 | 316 | Montevideo | SRR8272067 | 2018 | G |
| OSF088277 | USA:CA | Nuts | PDS000027237 | 316 | Montevideo | SRR8274725 | 2018 | G |
| OSF088278 | USA:CA | Nuts | PDS000027237 | 316 | Montevideo | SRR8274723 | 2018 | G |
| OSF088279 | USA:CA | Nuts | PDS000027237 | 316 | Montevideo | SRR8272153 | 2018 | G |
| OSF088282 | USA:CA | Nuts | PDS000027237 | 316 | Montevideo | SRR8272155 | 2018 | G |
| OSF088283 | USA:CA | Nuts | PDS000027237 | 316 | Montevideo | SRR8272614 | 2018 | G |
| PNUSAS001586 | USA | Clinical | PDS000027237 | 316 | Montevideo | SRR3277620 | Missing |  |
| PNUSAS001648 | USA | Clinical | PDS000027237 | 316 | Montevideo | SRR3277288 | 2016 |  |
| PNUSAS001685 | USA | Clinical | PDS000027237 | 316 | Montevideo | SRR3270925 | 2016 |  |
| PNUSAS001723 | USA | Clinical | PDS000027237 | 316 | Montevideo | SRR3289850 | 2016 |  |
| PNUSAS001724 | USA | Clinical | PDS000027237 | 316 | Montevideo | SRR3270998 | 2016 |  |
| PNUSAS001745 | USA | Clinical | PDS000027237 | 316 | Montevideo | SRR3495148 | Missing |  |
| PNUSAS001799 | USA | Clinical | PDS000027237 | 316 | Montevideo | SRR3289840 | 2016 |  |
| PNUSAS001966 | USA | Clinical | PDS000027237 | 316 | Montevideo | SRR3473884 | 2016 |  |
| PNUSAS002790 | USA | Clinical | PDS000027237 | 316 | Montevideo | SRR3897960 | 2016 |  |
| PNUSAS004962 | USA | Clinical | PDS000027237 | 316 | Montevideo | SRR5031400 | 2016 |  |
| PNUSAS008631 | USA | Clinical | PDS000027237 | 316 | Montevideo | SRR5336282 | Missing |  |
| PNUSAS017345 | USA | Clinical | PDS000027237 | 316 | Montevideo | SRR5864604 | Missing |  |
| PNUSAS034916 | USA | Clinical | PDS000027237 | 316 | Montevideo | SRR6870271 | 2018 |  |
| PNUSAS041766 | USA | Clinical | PDS000027237 | 316 | Montevideo | SRR7235831 | Missing |  |
| PNUSAS045784 | USA | Clinical | PDS000027237 | 316 | Montevideo | SRR7633156 | Missing |  |
| PNUSAS049251 | USA | Clinical | PDS000027237 | 316 | Montevideo | SRR7688270 | Missing |  |
| PNUSAS051029 | USA | Clinical | PDS000027237 | 316 | Montevideo | SRR7739852 | Missing |  |
| PNUSAS136046 | USA | Clinical | PDS000027237 | 316 | Montevideo | SRR11174347 | Missing |  |
| 12-1128 | Canada | Clinical | PDS000032600 | 138 | Montevideo | SRR5055291 | Missing |  |
| CFSAN010209 | USA:CA | Pistachio | PDS000032600 | 138 | Montevideo | SRR5384655 | 2009 | A |
| FSIS31800836 | USA:UT | Beef | PDS000032600 | 138 | Montevideo | SRR7657490 | 2018 |  |
| FSLR9-1449 | USA:NY | Clinical | PDS000032600 | 138 | Montevideo | SRR8502795 | 2013 |  |
| OSF005645 | USA:CA | Pistachio | PDS000032600 | 138 | Montevideo | SRR958039 | 2009 | G |
| OSF056829 | USA:CA | Raw Pistachios | PDS000032600 | 138 | Montevideo | SRR5120042 | 2014 | I |
| OSF067822 | USA:CA | Environmental sponge | PDS000032600 | 138 | Montevideo | SRR5991129 | 2017 | G |
| OSF069191 | USA:CA | Environmental sponge | PDS000032600 | 138 | Montevideo | SRR6109466 | 2015 | G |
| PNUSAS004495 | USA:WY | Cattle | PDS000032600 | 138 | Montevideo | SRR4418296 | Missing |  |
| PNUSAS028544 | USA | Clinical | PDS000032600 | 138 | Montevideo | SRR6317156 | Missing |  |
| PNUSAS044191 | USA | Clinical | PDS000032600 | 138 | Montevideo | SRR7429448 | Missing |  |
| PNUSAS072017 | USA:MD | Clinical | PDS000032600 | 138 | Montevideo | SRR9972052 | Missing |  |
| PNUSAS132560 | USA | Clinical | PDS000032600 | 138 | Montevideo | SRR11005270 | Missing |  |

### In silico multi-locus sequence typing (MLST)

Initial sequence analysis of the isolates was performed using an *in silico* MLST approach, based on information available at the *Salmonella enterica* MLST database (25). Seven loci (*aroC, dnaN, hemD, hisD, purE, sucA,* and *thrA)* were used for MLST analysis to generate a sequence type (ST).

### Genome Closure

Five *Salmonella* Senftenberg genomes and five *Salmonella* Montevideo genomes isolated from pistachios were completely closed using long read sequencing technology for this study (Table 2). Six of the *Salmonella* isolates were sequenced with the PacBio Sequel system (Pacific Biosciences, Menlo Park, CA, USA) and four isolates were sequenced with the Oxford Nanopore GridION (Oxford Nanopore Technologies, Oxford, UK). The ten closed genomes were submitted in DDBJ/EMBL/GenBank, and annotated using the NCBI Prokaryotic Genome Annotation Pipeline (PGAP) (26). Eight of the complete genomes were reported with a detailed method description in two separate genome announcements (27, 28).

**Table 2:** Closed genome statistics from this study.

| Sample ID | Serovar | MLST ST | Collection Year | Chromosome/<br>Plasmids Size (bp) | Sequencing Technology | Coverage | GenBank Accessions |
| --- | --- | --- | --- | --- | --- | --- | --- |
| CFSAN000258 | Montevideo | ST316 | 2009 | 4,694,397 | PacBio | 500x | CP029035 |
| CFSAN045764 | Montevideo | ST316 | 2016 | 4,694,399 | PacBio | 375x | CP029039<sup>(27)</sup> |
| CFSAN051296 | Montevideo | ST316 | 2016 | 4,694,373 | PacBio | 204x | CP029336<sup>(27)</sup> |
| CFSAN005645 | Montevideo | ST138 | 2009 | 4,615,193 | ONT | 103x | CP040380<sup>(25)</sup> |
| CFSAN010209 | Montevideo | ST138 | 2009 | 4,619,529 | ONT | 73x | CP040379<sup>(25)</sup> |
| CFSAN000878<br>pCFSAN000878 | Senftenberg | ST14 | 2009 | 4,794,849<br>319,930 | PacBio | 479x | CP029036<br>CP029037 |
| CFSAN045763 | Senftenberg | ST14 | 2016 | 4,766,139 | PacBio | 275x | CP029038<sup>(27)</sup> |
| CFSAN047866 | Senftenberg | ST14 | 2016 | 4,807,052 | PacBio | 477x | CP029040<sup>(27)</sup> |
| CFSAN010508 | Senftenberg | ST185 | 2013 | 4,820,913 | ONT | 77x | CP037894<sup>(25)</sup> |
| CFSAN087304 | Senftenberg | ST185 | 2018 | 4,819,673 | ONT | 25x | CP037892<sup>(25)</sup> |

A brief description of the methods for Pacbio sequencing is as follows. Genomic DNA was isolated using the DNeasy Blood and Tissue kit (Qiagen, Inc., Valencia, CA, USA), and libraries were prepared following the 20-kb PacBio sample preparation protocol. Genomes were sequenced using a single-molecule real-time (SMRT) cell with the PacBio Sequel sequencing system, following the manufacturer’s specifications. Using the Hierarchical Genome Assembly Process version 4.0 (HGAPv4) tool with default parameters, the genomes were *de novo* assembled and resulted in a single chromosome for each of the isolates and one plasmid (pCFSAN000878).

For MinION sequencing, the genomic DNA was isolated using the Maxwell RSC Cultured Cells DNA kit (Promega, Madison, WI). Extraction was performed according to manufacturer’s recommendations with the addition of RNase A after lysis. Libraries were prepared using the Oxford Nanopore Rapid Barcoding kit (SQK-RBK004) and run using a FLO-MIN106 (R.9.4.1) flow cell on a GridION instrument, according to the manufacturer’s instructions for 48 hours. The reads were assembled following previously reported analysis workflows (29). Briefly, the nanopore-generated raw reads were *de novo* assembled using Canu v1.7 (30). A second polished assembly was generated using SPAdes hybrid assembly (24) (with default settings) using both the Nanopore-generated raw data and the Miseq-generated raw data for each isolate. The two assemblies were then compared for synteny using Mauve aligner version 20150226 (31) and if in agreement the polished assembly was submitted to NCBI. The *de novo* assembly resulted in a single chromosome for each of the four isolates.

### Gene-by-gene comparison approach

To provide gene-by-gene comparison, we aligned all of our closed genomes based on serovar using MAUVE aligner version 20150226 using the progressive algorithm with default settings, and phages were identified using PHASTER (32). Any unique genomic regions, phages, and plasmids identified were imported into Ridom SeqSphere^+^ v. 6.0.2 (Ridom GmbH, Munster, Germany) (33) and used as a reference to interrogate the 201 short read isolate assemblies used in this study for presence or absence.

### Whole genome MLST (wgMLST) analysis

Targeted core genome MLST (cgMLST) and accessory genome analysis of the isolates associated with pistachio outbreaks and recalls were performed using Ridom SeqSphere^+^ software v 6.0.2 (33) by creating *ad hoc* wgMLST databases derived from the closed genomes previously described. Using the cgMLST target definer tool and accessing our genomes from NCBI, an *ad hoc* wgMLST scheme for *Salmonella* Senftenberg was developed. The scheme was based on the closed reference genome CFSAN047866 (CP029040), comprised of 4,768 genes, and query genomes (CP029036, CP029038, CP037892, and CP037894) found in Table 2 using the following default thresholds. For the reference genome filter thresholds: (1) minimum length: 50 bases; (2) start codon and a single stop codon required; (3) homologous gene filter, excluding multiple copies of a gene with identity ≥ 90% or more than 100 bases overlap; (4) overlap filter, if overlap > 4 bases. For the query genome filters: (1) 90% gene identity and 100% gene overlap in all query genomes; (2) BLAST options: word size = 11, mismatch penalty = −1, match reward = 1; gap open costs = 5, gap extension costs =2. The resulting scheme consists of 4,548 loci (3,908 core targets and 640 accessory targets). The 106 *Salmonella* Senftenberg assemblies (54 nuts/seeds, 44 environmental, 7 clinical and 1 poultry) were typed using this scheme. The alleles identified were used to establish a matrix disregarding missing allele values. A minimum spanning tree was constructed to visualize the results using the Ridom SeqSphere^+^ software. Clusters were automatically assigned using the default distant threshold for the software of ≤ 10 alleles.

The same process was repeated to create an *ad hoc* wgMLST scheme for *Salmonella* Montevideo using the closed genome CFSAN051296 (CP029336), comprised of 4,664 genes, as the reference and the four other closed genomes from this study (CP029035, CP029039, CP040379, and CP040380) as query genomes using the previously defined settings to define the core genome loci. The resulting scheme consists of 4,447 loci (3,876 core targets and 571 accessory targets). The 95 *Salmonella* Montevideo assemblies (49 nuts/seeds, 30 clinical, 9 environmental, 2 poultry, 2 missing source, 1 beef, 1 swine, and 1 cattle) were typed using this scheme. A minimum spanning tree of all isolates was constructed using the software from the allele calls disregarding missing values, and clusters were defined with the same threshold as above.

### Phylogenetic SNP analysis using CFSAN SNP Pipeline

Due to the large allele differences identified in the wgMLST analysis, phylogenetic SNP analysis was performed separately on the four different sequence types of *Salmonella* Senftenberg and Montevideo using the CFSAN SNP Pipeline (34). In short, WGS reads were mapped to the reference genome using Bowtie2 and the resulting files were processed using SAMtools. The variant sites were identified using VarScan, and a SNP matrix was produced using custom scripts. Four SNP matrices were generated using the closed genomes CFSAN000878 (ST14), FSW0104 (ST185), CFSAN000258 (ST316), and CFSAN005645 (ST138) as references. The SNP matrices can be found in Tables S1-S4. Phylogenetic trees were constructed using the RAxML (Randomized Accelerated Maximum Likelihood) program with 500 bootstrap replicates (35). All RAxML analyses were performed with the default parameter settings and the GTRCAT nucleotide substitution model. The trees were visualized using Figtree software (version 1.4.4).

### Pistachio storage study

A storage study was conducted to evaluate the ability for *Salmonella* (including *Salmonella* Senftenberg and Montevideo isolates associated with the 2016 outbreak) to persist on raw, inshell pistachios in conditions simulating a range of typical storage conditions. A cocktail containing four different *Salmonella enterica* serovars (Anatum, Montevideo, Oranienberg, and Senftenberg) associated with low moisture food commodities and one strain of *Salmonella* Newport isolated from tomatoes was used in this experiment (Table 3). The genomes of all five isolates were completely closed using Pacbio long read technology (27, 36). For this study, pistachios were inoculated with the 5-strain cocktail, as well as inoculated with the same strains on an individual basis. Inocula were prepared from stock culture as described in Keller et al. (37) with slight modifications. Briefly, a single colony from each working stock was transferred to Trypticase Soy Broth (TSB, BD, Franklin Lakes, NJ, USA) and incubated overnight at 37°C. After incubation, 100 µl of inoculum was spread plated on Trypticase Soy Agar (TSA, BD) plates. The plates were incubated overnight at 37°C and then bacteria were harvested by adding 1.0 ml of Buffered Peptone Water (BPW, BD) to the surface of the plate and gently scraped using an L-shaped plate spreader. Each plate yielded ∼0.5 ml of harvested culture at approximately 10 log CFU/ ml. The cocktail was formed by combining equal volumes of all five harvested cultures and used for inoculation of the pistachios. Organic, raw inshell pistachios were purchased and tested for background microbial populations using the Biomerieux Tempo AC kit (Biomerieux, Durham, NC, USA). A high-population cell suspension was prepared by combining 18 ml of the five-serotype *Salmonella* cocktail with 1,800 ml of sterile deionized water. A total of 900 g of pistachios were soaked in the suspension bath for 1 minute, and then placed on trays lined with absorbent towels in a biosafety cabinet to airdry overnight at room temperature. After 24 h, pistachios were stored in desiccator cabinets equilibrated to either 35% relative humidity (RH) or 54% RH at 25°C, which were selected to represent a low and high water activity (a_w_) within the range of a_w_ levels typical of low moisture foods (38). Uninoculated control pistachios were held under the same conditions. To determine the surviving populations, triplicate (10g) samples were removed for analysis at 0, 1, 2, 4, 6, 13, 27, 55, 83, 180, 270, and 365 days. The water activity was recorded for the inoculated and control pistachios from the different relative humidity conditions using an Aqualab model 4TE water meter (Decagon Devices, Pullman, WA). Two random 10 g samples from the uninoculated control group were tested using the Biomerieux Tempo AC kit to determine background microbe counts. The triplicate samples of inoculated pistachios were diluted 1:9 in BPW and appropriate dilutions were plated in duplicate on the differential media m-TSAYE [Trypticase Soy Agar with Yeast Extract supplemented with 0.05% (wt/vol) ammonium iron citrate and 0.03% sodium thiosulfate]. After overnight incubation at 37°C, colonies were counted and log CFU/g was calculated. A repeated measure analysis of variance (ANOVA) was performed in R v3.6.1 to determine the differences in log CFU/g over time between the high and low humidity conditions. The differences in the slopes (or rate of reduction in log CFU/g) between the two humidity conditions were also investigated in R.

**Table 3:** Strains used in storage study to assess persistence in pistachios.

| Strain | Serovar | Source | MLST (ST) |
| --- | --- | --- | --- |
| CFSAN076215 | Anatum | Peanuts | 64 |
| CFSAN051296 | Montevideo | Pistachio | 316 |
| CFSAN000836 | Newport | Tomato | 118 |
| CFSAN076211 | Oranienburg | Pecans | 3613 |
| CFSAN045763 | Senftenberg | Pistachio | 14 |

Two methods were used to determine composition of surviving serotypes at different time points, the Luminex xMAP® *Salmonella* Serotyping Assay (Luminex, Madison, WI, USA) and metagenomic sequencing using the Illumina MiSeq benchtop sequencer (Illumina, San Diego, CA, USA). For the Luminex assay, 12 – 16 individual colonies from each condition were selected from the m-TSAYE plates and DNA was extracted using Bio-Rad InstaGene^TM^ matrix (Bio-Rad, Hercules, CA) following the manufacturer’s protocol for DNA extraction from bacterial cells. DNA was analyzed for molecular determination of serotype in *Salmonella* following the protocol published by the CDC (39, 40). For the sequencing assay, the initial 1:9 dilution of pistachio:BPW was incubated at 37°C overnight. A 2 ml aliquot of the overnight enrichment was centrifuged at 6000 RPM for 10 minutes and then DNA was extracted using the Maxwell RSC Cultured Cells DNA Kit (Promega, Madison, WI, USA). Extraction was performed according to manufacturer’s recommendations with the addition of RNase A after lysis. Sequencing libraries were prepared using Nextera DNA Prep Library kit (Illumina, San Diego, CA) following the manufacturer’s protocols. The finished libraries were sequenced on the Illumina MiSeq using v3 600-cycle sequencing chemistry with 2 × 250 paired end reads. Raw reads from the triplicate samples were concatenated to increase our limit of detection and then analyzed using the Short Read Sequence Typer Tool v2 (SRST2) (41) with the maximum number of mismatches per read set to 0, the maximum number of mismatches for reporting ST profile set to 0, and minimum mean percent coverage set at 80%. Since we were looking at the presence or absence of known serovars in our inoculum, the threshold for minimum mean percent coverage was lowered below 90%. This tool utilizes MLST to assign reads to alleles and determines STs, which allowed us to determine the serovar present, since our isolates have distinct STs.

The single-strain storage study was conducted in parallel to the 5-strain cocktail storage study. The same procedure was used to prepare a high population cell inoculum of each individual strain and to inoculate 400 g portions of pistachios with each serovar. The same storage conditions were used and triplicate (10 g) samples were removed for analysis at Day 0, 1, 7, 28, 56, and 84. The triplicate samples were diluted 1:9 in BPW and appropriate dilutions were plated in duplicate on the m-TSAYE for direct plate counts.

## Results and discussion

### In silico MLST

*In silico* seven locus MLST was performed on the 106 *Salmonella* Senftenberg and 95 Montevideo isolates identified from the Pathogen Detection browser. MLST results for the *Salmonella* Senftenberg isolates identified two different STs. The 54 isolates from NCBI SNP cluster PDS000031814 belong to ST14, while the 52 isolates from SNP cluster PDS000031739 belong to ST185. The ST14 group includes seven of the clinical isolates and 39 isolates from pistachios with a date of collection ranging from 2009 to 2016. The ST185 group contains 12 isolates from pistachios collected between 2013 and 2018. There were no clinical isolates associated with this ST.

The MLST results for *Salmonella* Montevideo isolates also detected two STs. SNP cluster PDS000027237 is a member of ST316, while SNP cluster PDS000032600 belongs to ST138. The ST316 group contains a total of 82 isolates, of which 24 were clinical isolates and 33 were isolates from pistachio sources ranging in collection date from 2009 to 2018. ST316 has previously been described as the “outbreak” clade due to its linkage with numerous outbreak-associated plant products including tahini, sprouts, spices, as well as pistachios (42). The ST138 group of 13 isolates contains 6 clinical isolates and 3 isolates from pistachios. These sequencing typing results can be found in Table 1 and clearly indicate that pistachios have been contaminated by multiple strains of *Salmonella* Senftenberg and Montevideo.

### Closed Genomes

Long Read Sequencing technology was employed to generate 10 closed genomes for representatives of each ST to use as a reference for SNP analysis, as well as to identify the positions of mobile elements and assess genetic differences between STs. These specific isolates were selected as they were sourced from pistachios, readily available in our laboratory, and had different collection dates. In this study, we used two different types of long read sequencing technology, Pacific Biosciences (PacBio) and Oxford Nanopore Technology (ONT), to generate the closed genomes (27, 28). The *S.* Senftenberg isolates ranged from 4.7 – 4.8 mb in size while the *S.* Montevideo isolates were ∼ 4.6 mb. Those isolates sequenced with PacBio had genome coverage from 204 -500×, and those sequenced with ONT had 25 – 103× coverage. One plasmid, over 300 kb in size, was found associated with *Salmonella* Senftenberg CFSAN000878. The genome statistics for these ten isolates can be found in Table 2.

### Genome-wide comparisons

The content of the ten closed genomes of *Salmonella* consisting of 2-3 representatives from each sequence type were compared to each other, since many NGS analysis algorithms mask mobile elements such as phages, plasmids, and pathogenicity islands and which may lead to important characteristics of an isolate being overlooked. By comparing the complete and closed genomes from different isolates, we can find and identify genomic elements that a clonal strain may acquire or dispose of over time. The closed genomes were evaluated for conserved genes shared within the same sequence type, as well as between both serovars (Senftenberg and Montevideo), using the Mauve genome aligner, PHASTER, and BLAST. As expected, the isolates within the same sequence type showed higher similarity.

Interestingly, one unique region was identified that was found to be shared between the ST14 and ST316 isolates but missing in the ST185 and ST138 isolates. This unique region was determined to be the Copper Homeostasis and Silver Resistance Island (CHASRI) and is about 21 kb in size (43). Based on analysis using Ridom SeqSphere^+^, this island was confirmed in all of the short read sequence data of isolates associated with ST14 (N=54) and ST316 (N=82). The island is located on the chromosome in both serovars and contains the genes *pcoABCDEFGRS*, *cusABCFRS, silE*, and *silP*. The sequence of the CHASRI had a 100% identity intra-serovar and a 99.95% identity inter-serovar when the region was analyzed with BLAST (44).

The CHASRI has been identified previously in *Escherichia coli, Enterobacter cloacae, Klebsiella pneumoniae,* and *Salmonella enterica* (43). This island has previously been shown to increase the ability of a bacterium to survive in an environment with higher levels of copper and silver, as well as the ability to provide tolerance during the transition between aerobic and anaerobic environments, affording a fitness advantage for facultative anaerobes (43, 45). The use of copper as a growth promoter in swine and poultry has led to the selection of this island in *Salmonella* associated with these sources in other countries (46, 47). The use of copper as a fungicide and bactericide in agriculture has been reported for many years, and it is generally considered safe for both conventional and organic produce use (48, 49). Copper deficiency is reportedly common for pistachios and the use of copper in foliar and soil treatments has been recommended to the industry (50). Copper-containing foliar sprays may be applied at various times while pistachios are developing in the trees (51). The presence of the CHASRI in the pistachio-associated isolates may be an acquired adaptation by these specific salmonellae in response to copper stress in the growing environment.

The IncH12/IncH12A plasmid (CP029037) identified in *Salmonella* Senftenberg CFSAN000878, with a length of 319,930 bp, contains multiple heavy metal resistance genes which confer arsenic resistance (*arsB, arsC, arsH)*, mercury resistance (*merA, merC, merD, merP, merR, merT)*, and tellurium resistance (*terABCDEF*), as well as a copy of *silE*, *cusA/czcA,* and *cusS*. The plasmid also contains one antimicrobial resistance gene, MCR-9.1(52, 53), which confers Colistin resistance and is of significant public health importance. The plasmid sequences for the duplicated genes, *silE, cusA/czcA, and cusS,* differ from the same genes found in the chromosome with 91.2%, 93.7%, and 95.1% identity, respectively. This plasmid was only identified in one other isolate (CFSAN016599) from pistachio also collected in 2009. Since the plasmid has not been identified in any isolates after 2009, it is very likely that it was lost over time due to its large size and the genes on this plasmid not being necessary but rather redundant for increased fitness due to the incorporated CHASRI already in the chromosome.

Phages can encode factors that increase the virulence of *Salmonella* by enhancing adhesion, intracellular survival, and host entry (54). Phages were identified using PHASTER, and those sequences were then used as references in Ridom SeqSphere^+^ to interrogate the short-read assemblies. One intact phage, Salmon_SPN3UB_NC019545, was found in one of the closed genomes from ST14. Specifically, this phage was found in the closed genome of CFSAN047866 (CP029040) as well as in the short read data from 7 of 54 isolates by analysis with Ridom SeqSphere^+^. Interestingly, these isolates were not associated with a single group on the SNP tree. The two closed isolates from ST185 contain two phages: Aeromo_phiO18P_NC009542 and Salmon_118970_sal3_NC031940. Using Ridom SeqSphere^+^ to analyze the short-read data, the first phage was found in 52 of 52 isolates, and the second phage was found in 5 of 52 isolates (missing only from FSE0085 [SRR1212317]). All five Montevideo closed genomes, from both ST316 and ST138, contained a common intact phage, Salmon_Fels2_NC010463, identified in all 95 isolates. The three ST316 closed genomes contained an additional phage, Salmon_vB_SosS_Oslo _NC_018279, which was identified in the short-read data from the 81 of 82 isolates (missing from PNUSAS017345 [SRR5864604]). The two closed genomes from ST138 isolates have an additional intact phage, Escher_D108_NC_013594, which was identified using Ridom SeqSphere^+^ in the short-read sequence data from all 13 isolates belonging to ST138.

### Whole genome MLST-based analysis

Core genome multilocus sequence typing (cgMLST)-based methods have been previously employed for analysis of *Salmonella enterica* outbreaks and clusters (55, 56) and are less computationally expensive in comparison with SNP analysis. An advantage of using an *ad hoc* cgMLST approach is the ability to analyze isolates without the need for a reference sequence. In this study, two *ad hoc* wgMLST schemes were created for analysis of the *S.* Senftenberg and *S.* Montevideo isolates.

The *Salmonella* Senftenberg wgMLST scheme of 4,548 loci (3,908 core targets and 640 accessory targets) that was produced using the five closed *Salmonella* Senftenberg genomes isolated from pistachios was used to analyze the assemblies of the 106 Senftenberg isolates selected for our study. A minimum spanning tree produced by analysis of all common allele calls from both the core genome and accessory genome shows that Senftenberg isolates in ST14 are distinct from isolates in ST185 by a difference of 3,534 alleles (Fig. 1). This allele difference between ST14 and ST185 is meaningful and illustrates that the primary contamination sources for these isolates are not the same. The analysis separated the isolates belonging to ST14 into three clusters (Clusters 1, 2, and 3) and the ST185 isolates into one cluster (Cluster 4). The pistachio isolates from ST14 have more branching and allele difference (range of 0-11) over an eight-year time period (2009–2017). The majority of the isolates belong to Cluster 1, which contains isolates associated with the 2016 outbreak. One pistachio isolate, CFSAN015119, from 2009 located in Cluster 2 had a higher allele difference (≥10) from the other pistachio isolates. Cluster 3 is located 20 alleles away from the nearest pistachio isolate and is associated with isolates from chicken. The pistachio-associated isolates from ST185 have low allele differences (0-7) over a five-year time period (2013-2018), which illustrates genome conservation and clonality. It is noteworthy that a distinct central node was found with the ST185 isolates. Finally, the isolate FSE0085, isolated from contaminated tahini, has 50 allele differences from the central node and is not related to Cluster 4.

**Figure 1:**
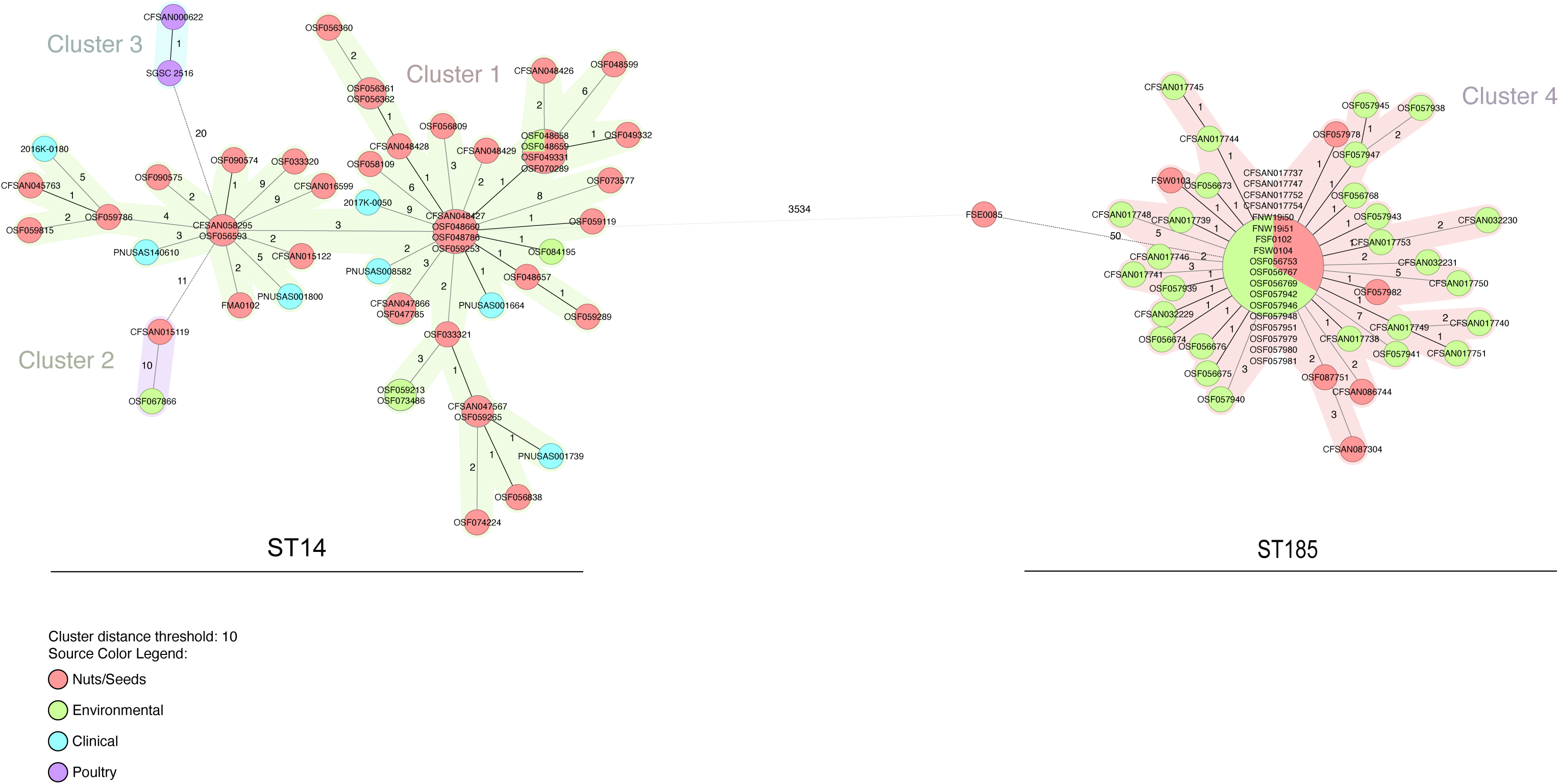
Minimum spanning tree of the wgMLST allelic profiles for *Salmonella* Senftenberg isolates (n = 106) using Ridom SeqSphere^+^. Distance based on wgMLST typing of 3,891 common loci (3,591 core genes and 300 accessory genes) using the parameters “pairwise ignoring missing values” during calculation. Isolates with 0 allele difference are found in the same node. Allele distances between samples are indicated, clusters are defined with maximum 10 alleles distance. Samples are color coded by their isolate source as in the legend.

The wgMLST scheme (3,876 core loci and 571 accessory loci) that was created using the five closed *Salmonella* Montevideo genomes isolated from pistachios was used to analyze the 95 *Salmonella* Montevideo isolates from our study. A minimum spanning tree produced by distance calculation for all common allele calls from both the core genome and accessory genome was constructed (Fig. 2), The wgMLST results show a distinct difference of 2,428 alleles between the isolates from ST316 and ST138. Similar to Senftenberg isolates, these isolates also show a sizeable allele disparity between the sequence types, which would be indicative of different sources of the contamination. Four well-defined clusters were identified, two for each ST (ST316 – Cluster 1 and 2, ST138 – Cluster 3 and 4). The isolates associated with the 2016 outbreak can be found in Cluster 1, while the majority of the isolates from pistachios are located in a single cluster for each ST (Cluster 1 and Cluster 3). A single isolate from trail mix (CFSAN015114 – 19 alleles) collected in 2008, as well as two clinical isolates (PNUSAS034916 – 47 alleles and PNUSAS051029 – 14 alleles) were found to fall outside of Cluster 1. Cluster 2 is associated with the chicken and swine isolates and is 18 alleles away from Cluster 1. The ST138 pistachio isolates are found in Cluster 3. Within this cluster, a pistachio isolate from 2009 (OSF005645) is a match and close relative of two isolates OSF067822 (environmental swab) and OSF056829 (pistachios) collected in 2014. This observation points to substantial conservation within the genome over a 5 year period.

**Figure 2:**
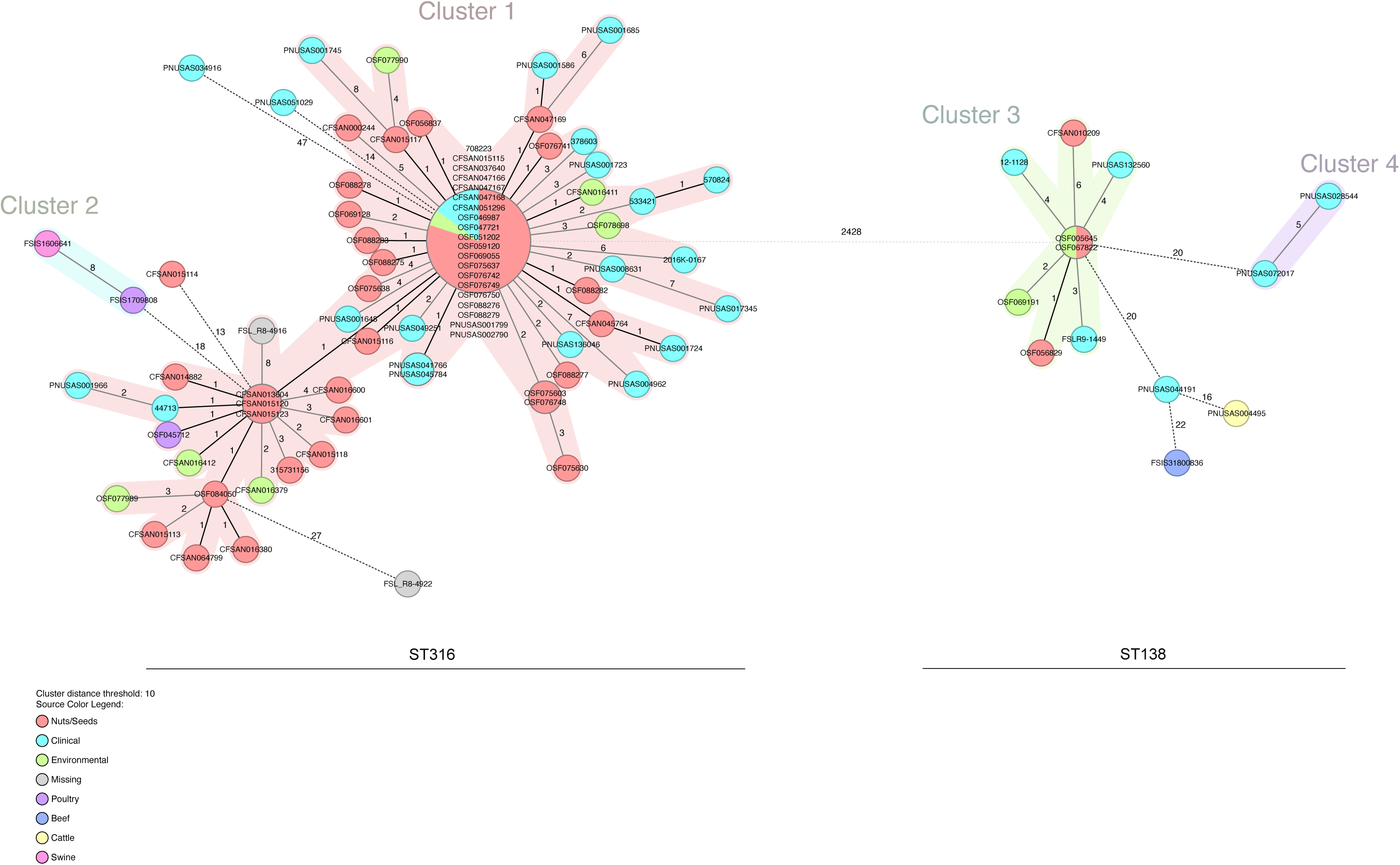
Minimum spanning tree of the wgMLST allelic profiles for *Salmonella* Montevideo isolates (n = 95) using Ridom SeqSphere^+^. Distance based on wgMLST typing of 3,876 common loci (3,634 core genes and 242 accessory genes) using the parameters “pairwise ignoring missing values” during calculation. Isolates with 0 allele difference are found in the same node. Allele distances between samples are indicated, clusters are defined with maximum 10 alleles distance. Samples are color coded by their isolate source as in the legend.

Interestingly, five clinical isolates from the United Kingdom were found in proximity to the outbreak Cluster 1 of pistachio isolates (0 – 4 alleles) (Fig. 2). One of these isolates was collected prior to the outbreak in 2014, and four were collected after the outbreak (2017-2019). There are two clinicals, PNUSAS041766 and PNUSAS045784, that are a match to each other with our wgMLST scheme and are one allele from the large node of outbreak-associated isolates. Also, three clinical isolates with 3-4 allele differences from an ST138 pistachio isolate are in Cluster 3. One of these isolates, 12-1128, is from Canada. Unfortunately in this case, without epidemiological data and more complete metadata from these isolates, it is impossible to confirm association with outbreak strains or the consumption of contaminated pistachios.

The results of our wgMLST analysis also showed the presence of large differences between isolates within the same serovar but belonging to different STs. For *Salmonella* Senftenberg, we were able to visualize the genomic separation of the two ST clusters, witness multiple mutation events in the ST14 strains, and see clonality within the ST185 isolates. The results from *Salmonella* Montevideo also showed the delineation of isolates from the different STs. We also identified multiple clinical isolates from the UK, Canada, and US that were not previously associated with an outbreak but were closely related to previous pistachio outbreak isolates from the US. The genomic connection of these closely related clinical isolates highlights the benefit of global partnerships in the area of food safety and public health and the need for increased metadata inclusion and transparency (57).

### SNP analysis of the four STs

To determine relatedness between reoccurring isolates, a reference-based SNP analysis was performed. The CFSAN SNP pipeline (34) is a reference-based variant detection analysis and is compatible for isolates that are closely related. The application should only be used on isolates that belong to the same NCBI SNP cluster (58). This pipeline was utilized here to produce four different SNP matrices using our closed genomes, representatives from each ST, from our study as reference genomes.

Using CFSAN000878 as the reference for the reference-based SNP analysis of the 55 *Salmonella* Senftenberg isolates that are typed as ST14, 146 variant positions were identified and a maximum likelihood phylogenetic tree was constructed using RAxML (Figure 3a). Anonymized facility identifiers were used for those isolates where the source facility was known. Isolates related to the 2016 outbreak are in bold font. Clinical isolates are highlighted on the tree with a star (n=7), and those clinical isolates associated with the 2016 outbreak are shown with filled-in stars (n=2). The isolates CFSAN000622 and SGSC 2516, both isolated from chicken, were the nearest neighbors to the pistachio isolate clade, with a range of 26 – 40 SNP differences. The pistachio isolates from 2009 – 2017 share a common ancestor, as highlighted in Figure 3a, and do not form distinct lineages based on the year of isolation. The isolates, both clinical and from pistachio associated with the 2016 outbreak (marked in bold), have a median of 4 SNPs and a range of 0 – 9 SNPs. These isolates appear throughout the tree, suggesting that multiple subpopulations of environmental *Salmonella* strains are potentially contaminating pistachios.

**Figure 3:**
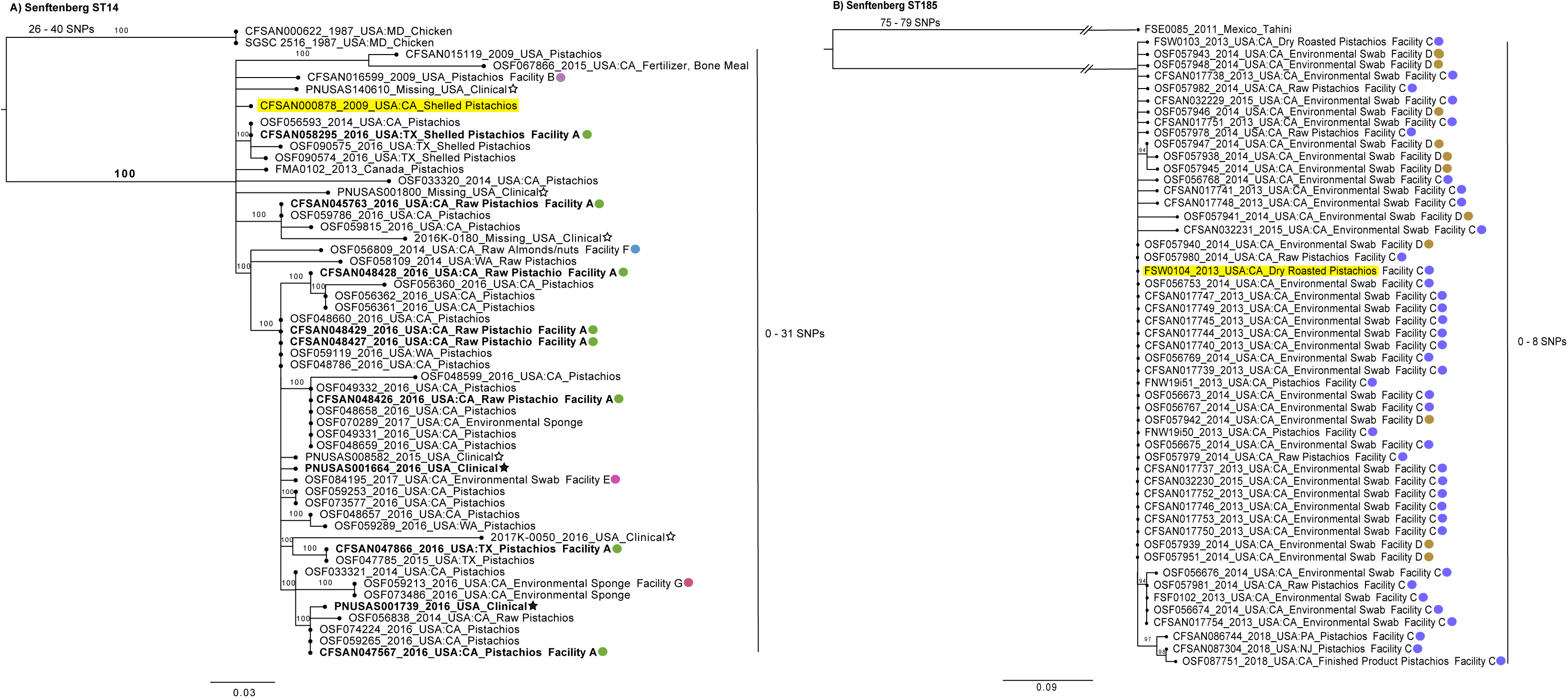
Phylogenetic analysis based on SNPs found in the *Salmonella* Senftenberg strains of this study. A SNP matrix was generated for both sets of isolates based on ST with the CFSAN SNP Pipeline (34). The SNP matrix was analyzed using RAxML using the GTRCAT substitution model and 500 bootstrap replicates. Reference strains are highlighted in yellow. Clinical isolates have a star symbol; black star = outbreak associated. Facility identifiers are also highlighted on the tree with different color circles. A) Maximum likelihood tree based on SNPs found in the 55 isolates from the ST14 group. B) Maximum likelihood tree based on SNPs found in the 52 isolates from the ST185 group.

Akin to this observation, five other clinical isolates that are not associated with the outbreak also have relatively close relationships to pistachio isolates. For instance, PNUSA008582 is 1 SNP away from both CFSAN048429 and CFSAN048427, while the other four clinicals have a minimum SNP distance of 5 – 13 SNPs from the nearest 2016 pistachio outbreak strain. The main clade has a total diversity range of 0 – 31 SNPs with a median of 7 SNPs. Previous research has shown SNP differences in *Salmonella* outbreaks to be <21 SNPs (18, 23, 59–61) with an outlier in serovar Agona with 30 SNP differences (62). From our study, a 2015 fertilizer (bone meal) isolate, OSF067866, was noted to be 9 SNPs distant from a 2009 pistachio isolate, CFSAN015119. While a definitive linkage of these two isolates is unattainable, it is notable that the use of pistachio hulls as a soil amendment has been explored (63–65). Additional examination of soil amendments and fertilizer blends containing pistachio hulls may shed additional light on potential reservoirs and persistence of *Salmonella* in orchard soil. The diversity of the strains based on the range of SNP differences in the ST14 cluster is indicative of *Salmonella* residence in pistachio primary production environment (i.e., orchard). Further evidence supporting *Salmonella* residence in this environment is that there are isolates collected from five other facilities (including pistachio and environmental sources), located in California and not linked to the 2016 outbreak, over multiple years found throughout the tree. Finally, the finding that all of these isolates contain the CHASRI island – coinciding with the known use of copper in pistachio orchards providing selective pressure for this adaptation, as well as previous studies showing poultry isolates containing this island – further supports that these strains have established residence in or around the production environment. Although, the initial source of the contamination (e.g., adjacent animal operations) of this strain can not be determined, our analysis shows that it was able to establish residence.

For the second Senftenberg tree, closed genome CFSAN010508 was used as the reference genome to analyze the 52 isolates identified by MLST to belong to ST185 (Figure 3b). The SNP analyses identified 108 variant positions within these isolates. The isolate FSE0085, tahini from Mexico, was the closest related isolate (75 – 79 SNPs) to the majority of the isolates (n=51) that grouped together to form a single monophyletic clade. The 51 isolates in this clade originated from pistachios and environmental swabs collected from two different facilities, Facility C and Facility D, both located in California. These isolates have not been associated with outbreaks but their isolation has resulted in product recalls. Very little variability exists between the isolates from these two facilities, with a median of 1 SNPs distance and a range of 0 – 8 SNPs over a five-year period. The isolates collected in 2018 form a group that is a median of 4 SNPs distant (range 3 – 8 SNPs) from those collected in 2013, 2014, and 2015. The low SNP difference is indicative of a persistent strain that is not under stress and experiencing slower than normal evolutionary events (66). Some theoretical explanations for the presence of *Salmonella* isolates from these two facilities with 0 SNP distances between them include *Salmonella* establishing residence in the facilities after both receiving pistachios contaminated by a possible shared grower, harvester, or transporter in or before 2013 or a business relationship that allows for pistachios or equipment to be shared or exchanged between the two facilities. It seems unlikely that two strains with no SNP differences would occur in disparate facilities without some type of indirect or direct relationship between them. Rather, the small number of genetic differences over an extended period of time would suggest the transmission of the same strain between the two facilities and therefore a clonal strain taking harborage (20, 67). Further, the lack of any measurable SNP distance between these *Salmonella* isolates from these two facilities suggests that such residence would be in an environment with relatively low environmental stress, such as an environment devoid of implementation of more robust sanitation controls.

A total of 107 distinct SNP variants were identified by reference-based SNP analysis in the isolates of ST316 using CFSAN000258 as the reference genome. A resultant phylogenetic tree, capturing the relationship among the 82 isolates belonging to ST316, yielded several important findings (Fig. 4a). Two isolates, FSIS1709808 and FSIS16006641, isolated from chicken and swine, respectively, were 23 – 35 SNPs away from the main clade of the tree. Isolates related to the 2016 outbreak are in bold font and are not associated with one specific lineage on the tree. Clinical isolates are highlighted with a star on the tree (n=21), and those clinical isolates associated with the 2016 outbreak are shown with filled-in stars (n=8). There are isolates from seven different facilities found on this tree. The paraphyletic grouping of the clade breaks down into smaller clusters, with a total SNP distance ranging from 0 – 16 SNPs with a median of 3 SNPs. The 2016 outbreak strains have a median distance of 2 SNPs (range 0 – 10 SNPs).

**Figure 4:**
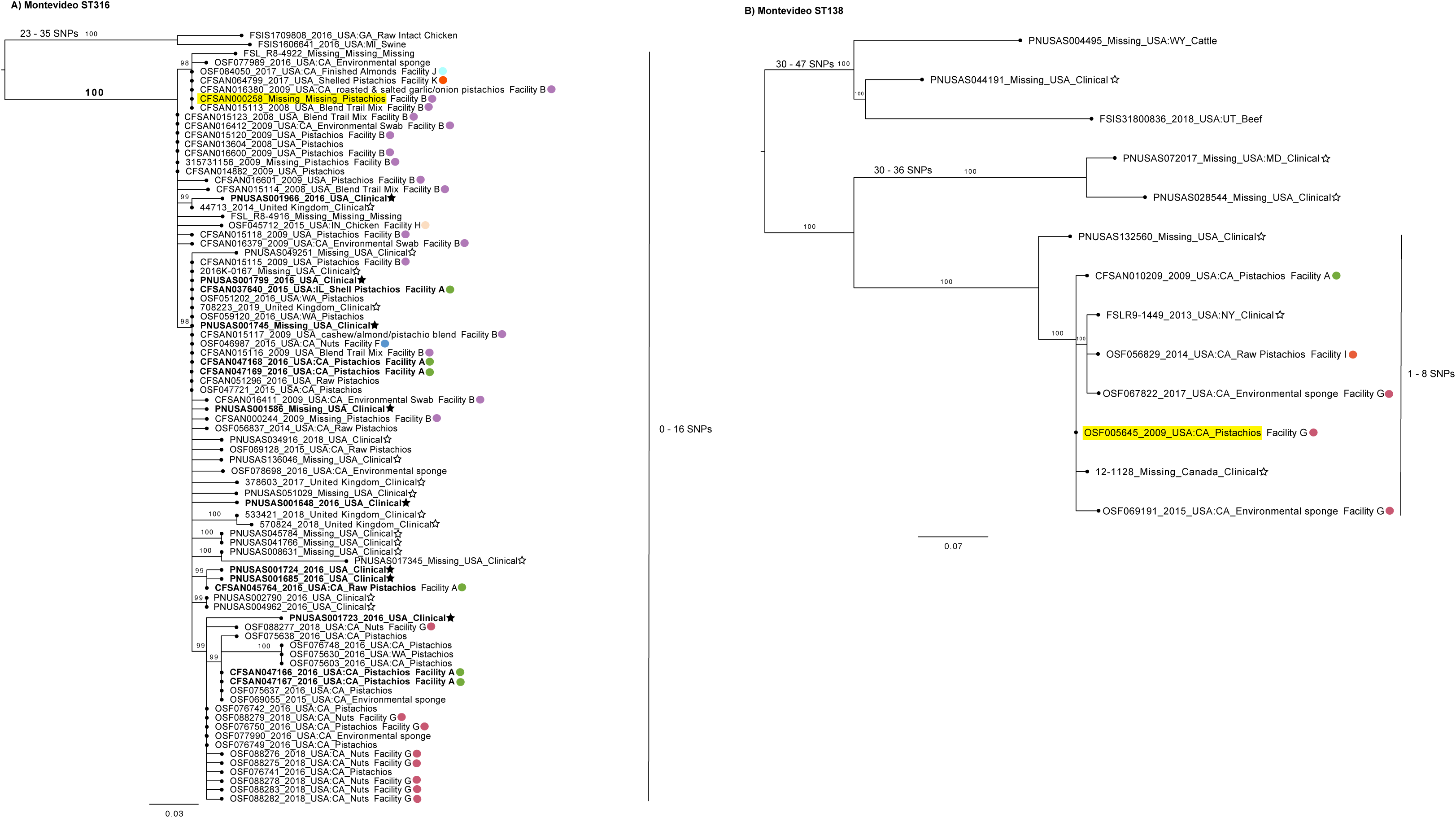
Phylogenetic analysis based on SNPs found in the *Salmonella* Montevideo strains of this study. A SNP matrix was generated for both sets of isolates based on ST with the CFSAN SNP Pipeline (34). The SNP matrix was analyzed using RAxML using the GTRCAT substitution model and 500 bootstrap replicates. Reference strains are highlighted in yellow. Clinical isolates have a star symbol; black star = outbreak associated. Facility identifiers are also highlighted on the tree with different color circles A) Maximum likelihood tree based on SNPs found in the 82 isolates from the ST316 group. B) Maximum likelihood tree based on SNPs found in the 24 isolates from the ST138 group.

Additionally, other clinical isolates from the United States and the United Kingdom were found to be closely related, with a minimum SNP distance range of 0 –11 SNPs and 0 – 4 SNPs, respectively. With the continued rise in imported/exported goods, the observation of the large number of clinical isolates that are clustered near pistachio isolates from both the United States and the United Kingdom further emphasizes the benefit of a genome-based globally connected surveillance system for foodborne pathogen. Isolates from seven different facilities collected over different points in time are detected throughout the tree. The presence of a chicken isolate (OSF045712) from Facility H located within the California nut cluster can possibly be explained by the use of a tree nut supplemented animal feed (68–70). The chicken isolate is 2 – 5 SNPs away from nuts in the cluster. This potential connection between a contaminated component of animal feed and livestock that eventually enters the food supply for humans illustrates the interconnectedness of many food production systems and the importance of a “One Health Approach” (71) for food safety. Similar to the Senftenberg ST14 group, the CHASRI is located in each of these isolates. This observation, together with the changes occurring over eight years (0-16 SNPS), are consistent with the possibility of a strain in the pistachio production environment that is most likely occurring in conjunction with *Salmonella* Senftenberg.

A phylogenetic tree was also constructed for the 13 Montevideo isolates from ST138 (Fig. 4b). A total of 48 distinct SNP variants were identified by reference-based SNP analysis in the isolates of ST138 using CFSAN010508 as the reference genome. Clinical isolates are highlighted with a star on the tree (n=6). This sequence type has been reported to be primarily associated with bovine sources (25), and indeed, two bovine isolates, PNSUAS004495 and FSIS31800836, were found to be distantly related by 30 – 47 SNPs from the pistachio isolates. There are three pistachio isolates from three different facilities (Facility A, Facility G, and Facility I). The cluster that includes the three pistachio isolates shows high relatedness with 1 – 4 SNPs (median 3 SNPs). Three isolates from Facility G, two environmental swabs and one pistachio, also belong to this cluster. The SNP distance between the strain isolated in 2009 and the 2015 isolate and 2017 isolate is only 2 SNPs. The isolates of this cluster, both pistachios and swabs from three different facilities (Facility A, Facility G and Facility I), were collected over a wide timespan (8 years) and have low diversity, which suggests a persistent *Salmonella* strain. This strain is likely to reside in a lower-stress environment than in the orchard environment, which could include shared farm equipment, transport vehicles, or an intermediate space. The low SNP distances suggest a strain that is not undergoing selective pressure and may also have a slower generation time similar to a resident pathogen (20). There are also three clinical isolates that are within 1 – 8 SNPs (median 3 SNPs) from the pistachio isolates. However, without epidemiological data we are unable to positively link these illnesses with contaminated pistachio exposure.

### Storage study to assess persistence in pistachios

In order to further explore the nature and likelihood of long-term environmental persistence of these *Salmonella*, we tested the persistence of *Salmonella* strains both individually and in a cocktail after inoculation on raw inshell pistachios. The strains used in this study were associated with low-moisture food products (Anatum-peanuts, Oranienburg-pecans, Montevideo-pistachios, Senftenberg-pistachios) and a strain isolated from tomato (Newport) (Table 3). Pistachios were inoculated with either a five-serovar cocktail of *Salmonella enterica* or one of the five *Salmonella enterica* strains and stored at one of two different relative humidities (35% RH and 54% RH) at 25°C. The Senftenberg and Montevideo strains used contain the CHASRI and are from ST14 and ST316, respectively (Table 3). The basis behind using the Newport strain was to determine if there were differences in survivability on a low-moisture food based on the initial source of the isolate (i.e., a high-moisture food). The *Salmonella* in the five-strain cocktail was able to persist in pistachios for up to one year (the length of the study) (Fig. 5). There was a 1 log CFU/g reduction in *Salmonella* during the period of desiccation (24 hour drying time) prior to storage. The *Salmonella* in the inoculated pistachios stored at 35% relative humidity (mean a_w_ = 0.3398, stdev = 0.0228) were able to persist for a year with only an additional 1 log CFU/g reduction after desiccation (a total of 2 log CFU/g reduction). The *Salmonella* in the pistachios stored at 54% relative humidity (mean a_w_ = 0.5337, stdev = 0.0370) showed an additional 3 log CFU/g reduction over a year, and a repeated measures ANOVA analysis showed the impact of humidity condition (high vs low) over time. The rate of reduction was higher for the high humidity condition based on the observed difference in the slope (rate of reduction in log CFU/g for low and high humidity was −0.124 and −0.264 per day, respectively) between the two conditions (*p*<.0001). Differences in log CFU/g between low and high humidity conditions was most pronounced after 6 months (Fig. 5). These findings are consistent with those reported in literature (37, 38).

**Figure 5:**
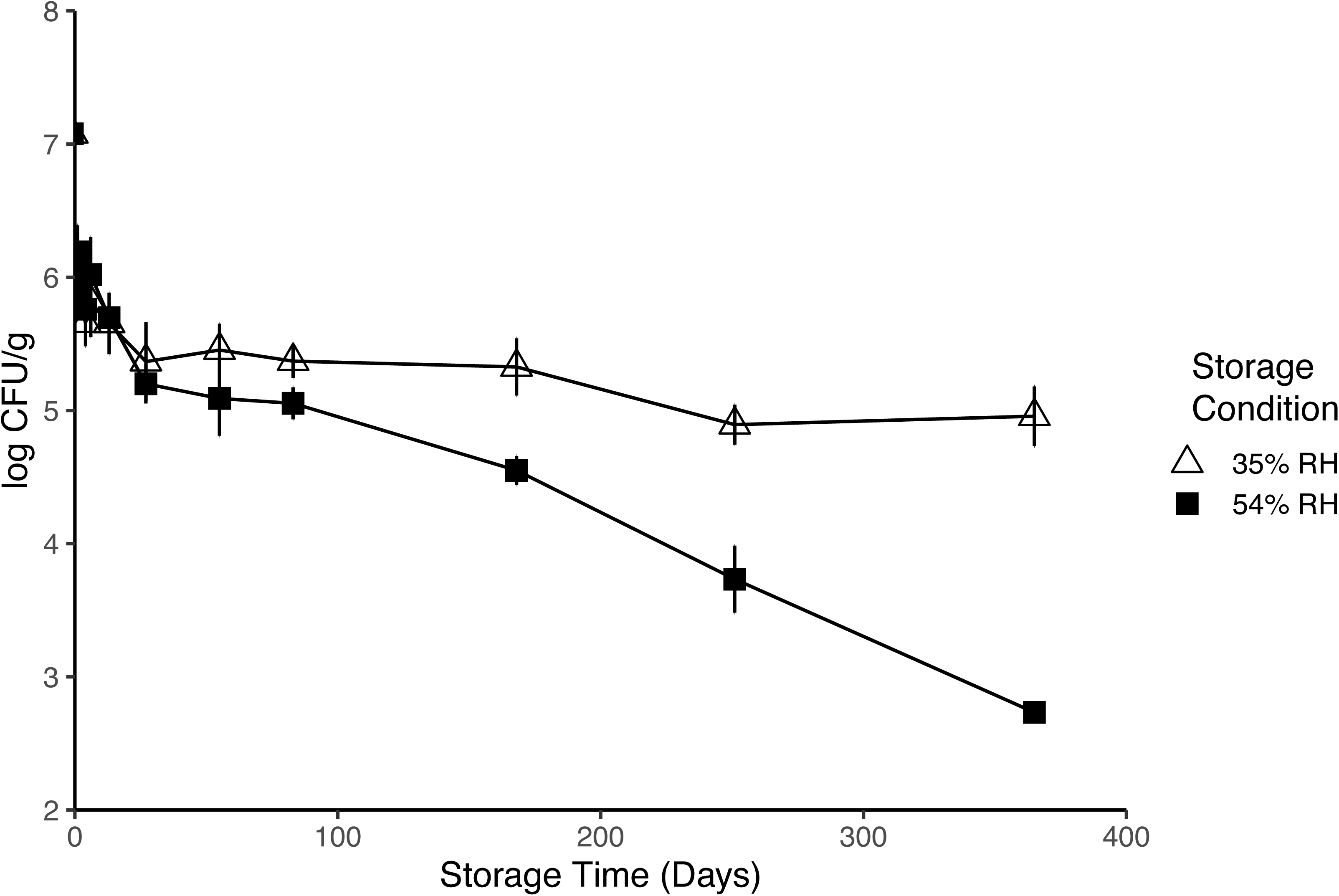
Survival curve of inoculated pistachios with a 5-strain *Salmonella* cocktail stored at different relative humidities. Measured *Salmonella* CFU/g values at different storage time points determined by direct plating methods from pistachios inoculated with a 5-strain cocktail of *Salmonella enterica*. The triangle represents the results from storage at 35% RH at 25°C and the black square represents the results from storage at 54% RH at 25°C. Each data-point is an average of at least 2 replicate measurements ± standard deviation.

One of our study aims was to determine if there was a difference in which serovars were surviving within the cocktail inoculum. A previous storage study by Kimber et al. (72) evaluated the surviving serovars from the cocktail at the end of the storage study, while we analyzed the composition of surviving serovars throughout the study, which allows for identification of when changes may have occurred. Two methods were used to determine the serovars present, the Luminex xMAP® *Salmonella* serotyping assay and metagenomic sequencing with data analysis using SRST2. The results of both methods are compared in Figure 6. Neither method correlates to relative abundance of the serovars in the inoculated pistachios but both methods provide a snapshot of what is surviving. The SRST2 analysis provides a more uniform picture of what serovars are present when compared to the molecular serotyping assay. Multiple instances were noted where colonies from a single serovar were not selected for the molecular serotyping assay but was able to be detected by metagenomic sequencing. Sequencing was conducted post-enrichment, therefore the serovars identified from the reads is from viable bacteria. The limitation of the molecular serotyping assay is the number of colonies that are selected and tested. Unintentional bias could be introduced into the results based on colony selection by either over- or under-selection, but there is a higher probability of sequencing any and all strains that are present above a certain threshold with the sequencing method. The combined serotyping results from both methods showed that all five serotypes could be detected at one year of storage. The results from the metagenomic ST assignment with SRST2 (Fig. 6b) concurred with the Luminex assay (Fig. 6a) results most of the time, but due to the limitation of sequencing depth, Oranienberg was not identified in several instances. The sequencing runs consisted of multiplexing 8-10 samples per run. However, this level of multiplexing did not provide enough sequencing depth to detect Oranienberg, which may have been occurring at lower concentrations. A subset of these samples were resequenced with 3-4 samples per run and detection of Oranienberg was possible for some of these repeats. Interestingly, the Kimber et al. 2012 study used the same Oranienburg strain as in our experiments, but the researchers did not identify this strain in the 100 colonies they tested at the end of their study (14.5 months) (72). In contrast, we were able to still recover the Oranienburg strain both in sequencing and by colony picks at 12 months. Two possibilities for the absence of Oranienburg in the Kimber et al. 2012 study could be that Oranienberg was missed due to the bias that can occur when picking colonies for serovar determination or Oranienberg simply did not survive. This could be determined by using metagenomic sequencing (removes colony pick bias) or by picking colonies at different time points (identify survival cutoff). The use of the metagenomic/bioinformatic pathway allows for a robust determination of the cocktail composition while reducing the bias and time of traditional molecular serotyping.

**Figure 6:**
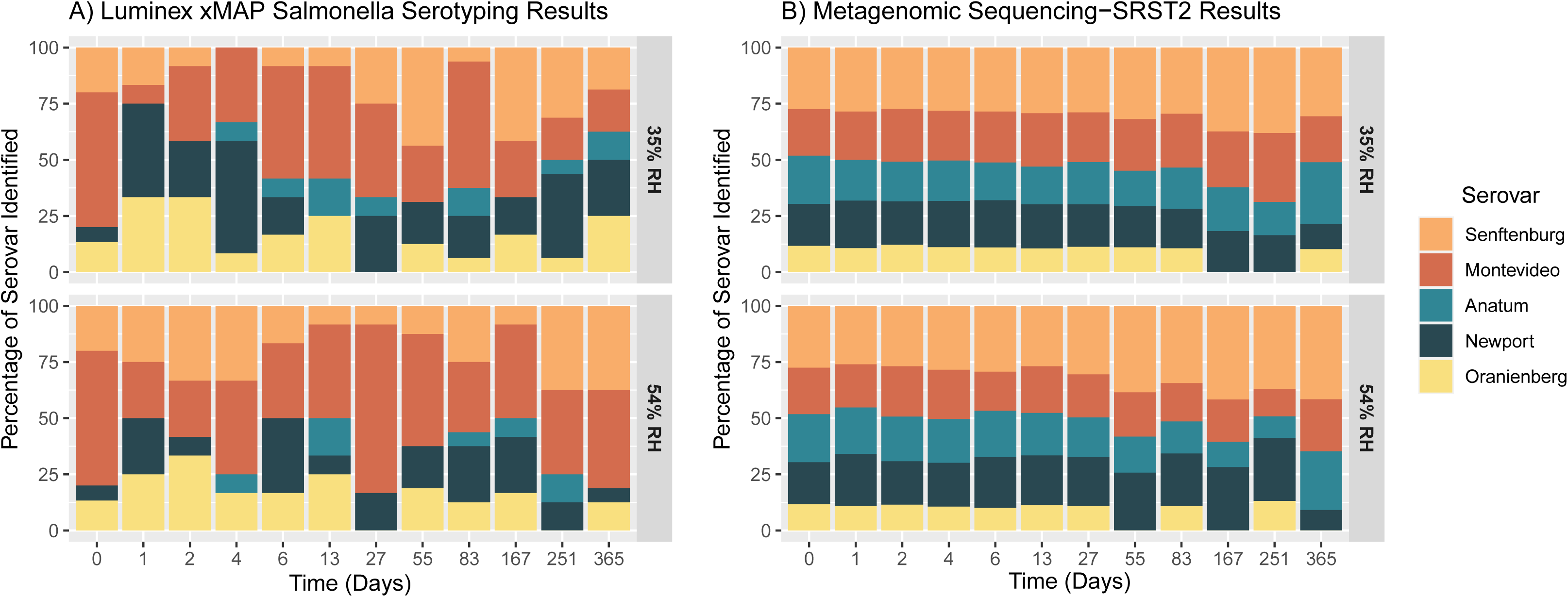
Comparison of two methods for serovar determination from the 5-strain cocktail inoculated pistachios. A) The results of the *Salmonella* molecular serotyping of colonies picked at different time points from inoculated pistachios stored at 35% and 54% RH. B) The results of the metagenomic sequencing of DNA isolated from overnight enrichment of inoculated pistachios at the same time points. The reads were analyzed using SRST2 with following parameters: maximum number of mismatches for reporting ST = 0; maximum number of mismatches per read = 0; and minimum mean percent coverage = 80%.

Each individual strain was also used to inoculate pistachios and stored at the same low and high relative humidity to compare the results of persistence of a single strain contamination versus a multi-strain contamination event. No significant differences in the bacterial counts between strains or between storage conditions (data not shown) were found during our three-month storage period, and therefore, these trials were halted at Day 84.

The results from our study conclude that the five-strain cocktail of *Salmonella* can survive in pistachios and be detected by direct plating methods when stored at different humidities for at least one year. There were no differences in the survivability of our strains when coexisting in a cocktail versus individually at three months. This storage study reinforces the capability for *Salmonella* to survive for extended times on this low-moisture commodity. Interestingly, the primary source of the *Salmonella* (low-moisture vs. high-moisture environment) may not necessarily determine the ability to survive in a different environment, shown with the survivability of the Newport strain on the pistachios as well. Taken together, the extended shelf-life of pistachios combined with our evidence of *Salmonella* survival for up to a year highlights the importance of long-term surveillance of this pathogen.

## Conclusions

Pistachios may be contaminated with pathogenic bacteria in the growing, production, and processing environments and present a hazard due to the ability for *Salmonella* to persist in low-moisture environments and foods. Diagnostic capabilities have been greatly enhanced due to the development and increasing applicability of WGS, which is now fully deployed as a molecular epidemiological tool to assist in foodborne outbreak investigations (17–19). Epidemiological data are an important complement to WGS for affirming exposure to a particular pathogen.

Nonetheless, by using WGS in two different phylogenetic analyses, we were able to match clinical and food *Salmonella* Senftenberg and Montevideo isolates and define SNP differences between closely related strains. Further, by implementing long read sequence technology, we identified the presence of the heavy metal tolerance island (CHASRI) in strains likely associated with preharvest contamination, indicative of their adaptation to the presence of copper in this niche. Presence of the CHASRI in *Salmonella* Senftenberg (ST14) and *Salmonella* Montevideo (ST316) may explain the increased level of recovery of these strains from pistachios or pistachio environments due to selective pressure from copper usage, even though previous data has shown a greater diversity of serovars present (10). Finally, this analysis points to at least three possible pistachio-associated environments where closely related *Salmonella* Senftenberg and Montevideo strains could have been residing as recently as 2018 including 1) multiple environmental adapted strains in the pre-harvest or orchard environment; 2) a clonal strain in the intermediate post-harvest environment (e.g., shared farm equipment or transport equipment); and 3) a clonal strain at the pistachio handling facilities, which could include storage silos.

Our storage study demonstrates that *Salmonella* will survive in pistachios for a minimum of one year at both high and low relative humidities. The use of a non-nut associated strain, *Salmonella* Newport, in our cocktail and its ability to be recovered at the 12 month time point, showed that the original source may not necessarily have a significant impact of survival of *Salmonella* on a low-moisture food. Pistachios may be stored untreated for 12 months or longer by pistachio handlers and then consumers may store them up to an additional 12 months (73). The ability of *Salmonella* to survive long-term in a dry environment highlights the need for adequate preventive controls in the processing of pistachios. Harris et al. (10) report that a majority of the pistachios sold in the U.S. market are roasted inshell. Identification and surveillance of the possible sources of contamination, (e.g., the growing environment, shared equipment, or processing environment) would likely augment implementation of specific processing changes that could further reduce the risk of contaminated nut products.

Taken together, this study shows the following: i) evidence of persistent *Salmonella* Senftenberg and Montevideo strains in pistachio environments as recently as 2018, ii) presence of the Copper Homeostasis and Silver Resistance Island (CHASRI) in *Salmonella* Senftenberg and Montevideo strains, suggesting an adaption in response to extrinsic pressures in the pistachio supply chain, and iii) use of metagenomic sequencing in conjunction with storage studies using a cocktail of *Salmonella* strains can identify strains that are persisting over time. This study further emphasizes the power of WGS in conjunction with metadata, which helps to define outbreaks and resident strains, as well as give insights into genetic factors that could lead to persistence in a given niche. Moreover, these findings underscore the importance of industry in adhering to food safety standards established for this commodity.

## Acknowledgments

The FDA Foods Program Intramural Funds supported the study.

